# Transcription regulation of SARS-CoV-2 receptor ACE2 by Sp1: a potential therapeutic target

**DOI:** 10.1101/2023.02.14.528496

**Authors:** Hui Han, Rong-Hua Luo, Xin-Yan Long, Li-Qiong Wang, Qian Zhu, Xin-Yue Tang, Rui Zhu, Yi-Cheng Ma, Yong-Tang Zheng, Cheng-Gang Zou

## Abstract

Angiotensin-converting enzyme 2 (ACE2) is a major cell entry receptor for severe acute respiratory syndrome coronavirus 2 (SARS-CoV-2). Induction of ACE2 expression may represent an effective tactic employed by SARS-CoV-2 to facilitate its own propagation. However, the regulatory mechanisms of ACE2 expression after viral infection remain largely unknown. By employing an array of 45 different luciferase reporters, we identify that the transcription factor Sp1 positively and HNF4α negatively regulate the expression of ACE2 at the transcriptional levels in HPAEpiC cells, a human lung epithelial cell line. SARS-CoV-2 infection promotes and inhibits the transcription activity of Sp1 and HNF4α, respectively. The PI3K/AKT signaling pathway, which is activated by SARS-CoV-2 infection, is a crucial node for induction of ACE2 expression by increasing Sp1 phosphorylation, an indicator of its activity, and reducing HNF4α nuclear location. Furthermore, we show that colchicine could inhibit the PI3K/AKT signaling pathway, thereby suppressing ACE2 expression. Inhibition of Sp1 by either its inhibitor mithramycin A or colchicine reduces viral replication and tissue injury in Syrian hamsters infected with SARS-CoV-2. In summary, our study uncovers a novel function of Sp1 in regulating ACE2 expression and suggests that Sp1 is a potential target to reduce SARS-CoV-2 infection.

## Introduction

Severe acute respiratory syndrome coronavirus 2 (SARS-CoV-2) is the causative pathogen of coronavirus disease 2019 (COVID-19) (Wu et al., 2020), which has spread rapidly across the world and caused a global public health event (Lu et al., 2020). The pathophysiological features of severe patients with COVID-19 are characterized by systemic inflammatory response, which manifests as acute respiratory distress syndrome and multiorgan dysfunction (Pan et al., 2021, van Eijk et al., 2021). SARS-CoV-2 cell entry depends on the SARS-CoV-2 receptor, angiotensin-converting enzyme 2 (ACE2), and the transmembrane serine protease, TMPRSS2 (Yan et al., 2020). During SARS-CoV-2 infection, the spike protein binds with ACE2 and undergoes cleavage by TMPRSS2 to allow the viral to fuse with host cell membrane, which is essential for viral entry (Hoffmann et al., 2020). ACE2 is wildly expressed in the lung, small intestine, kidney, liver, testis, heart and brain (Dong et al., 2020, Verdecchia et al., 2020). The expression of ACE2 in a wide variety of human tissues may explain why SARS-CoV-2 targets multiple organs.

Vaccination has been proven to be a highly effective way to prevent SARS-CoV-2 infection and reduce the incidence of hospitalization and death. However, due to the continuous mutation of the SARS-CoV-2, a number of individuals cannot mount an adequate immune response to COVID-19 through vaccination alone. The effectiveness of the different types of existing vaccines against the Omicron variants has been greatly reduced (Kuhlmann et al., 2022, Wilhelm et al., 2022). The spike protein of these SARS-CoV-2 variants shows an increased binding affinity to ACE2, leading to enhanced transmission ability (Liu et al., 2021, McCallum et al., 2022). Therefore, it is urgently needed to discover and develop a novel host-directed therapeutic against SARS-CoV-2, especially in aged population.

Accumulating evident indicates that SARS-CoV-2 infection significantly upregulates ACE2 expression (Gao et al., 2022, Wei et al., 2021, Xu et al., 2021a, Zhuang et al., 2020). SARS-CoV-2 infection upregulates the expression of HMGB1, which in turn induces ACE2 expression probably through an epigenetic mechanism (Wei et al., 2021). Furthermore, overexpression of SARS-CoV-2 spike protein significantly elicits ACE2 expression, which is dependent on the type I interferon signaling (Zhou et al., 2021). Indeed, ACE2 has been proved to be an interferon-stimulated gene (Ziegler et al., 2020). Meanwhile, two recent studies have demonstrated that androgen receptor positively regulates the expression of ACE2 at a transcriptional levels (Qiao et al., 2021, Samuel et al., 2020). Importantly, targeting the transcriptional regulation of ACE2 by reducing AR signaling, through androgen receptor antagonists or degraders attenuate SARS-CoV-2 infectivity (Qiao et al., 2021). A very recent study has demonstrated that ursodeoxycholic acid (UDCA), an inhibitor of the farnesoid X receptor (FXR), reduces ACE2 expression in human lung, intestinal, and liver organoids, thereby inhibiting SARS-CoV-2 infection (Brevini et al., 2022). Thus, ACE2 is a promising therapeutic target in the fight against COVID-19. However, the molecular mechanism underlying SARS-CoV-2 infection-induced ACE2 expression remains largely unknown.

To clarify the molecular mechanism underlying regulation of ACE2 by SARS-CoV-2 and colchicine, we employed a commercial array of 45 different luciferase reporters to assay a range of signaling pathways. Our data revealed that SARS-CoV-2 up-regulated ACE2 expression by activating the transcription factor Sp1 and inhibiting HNF4α through the PI3K/AKT pathway. Finally, we showed that inhibition of Sp1 by its inhibitor mithramycin A was active against SARS-CoV-2 in human respiratory epithelial cells and animal model. Thus, our findings suggest that Sp1 is an important transcription factor for ACE2 expression.

## Results

### SARS-CoV-2 infection up-regulates ACE2 expression, which is inhibited by colchicine treatment

As the receptor for the SARS-CoV-2 viral entry, ACE2 is regarded as a promising therapeutic target in the fight against COVID-19 (Monteil et al., 2020). Consistent with previous observations that SARS-CoV-2 infection significantly upregulates the mRNA expression of ACE2 (Gao et al., 2022, Wei et al., 2021, Xu et al., 2021a, Zhuang et al., 2020), we found that infection by SARS-CoV-2 upregulated the protein levels of ACE2 in HPAEpiC cells, a human lung epithelial cell line (Figure 1A and 1B). Similar results were obtained from immunofluorescence analysis (Figure 1C and 1D). Recent clinical studies have shown a mortality benefit with colchicine, a drug used for the treatment of acute gout and familial Mediterranean fever (Dasgeb et al., 2018, Gasparyan et al., 2015, Schlesinger et al., 2020, Slobodnick et al., 2015), when used in the treatment of COVID-19 patients (Drosos et al., 2022, Elshafei et al., 2021). In this study, we found that treatment with colchicine substantially inhibited ACE2 expression in HPAEpiC cells with or without SARS-CoV-2 infection (Figure 1A-D).

**Figure 1.**
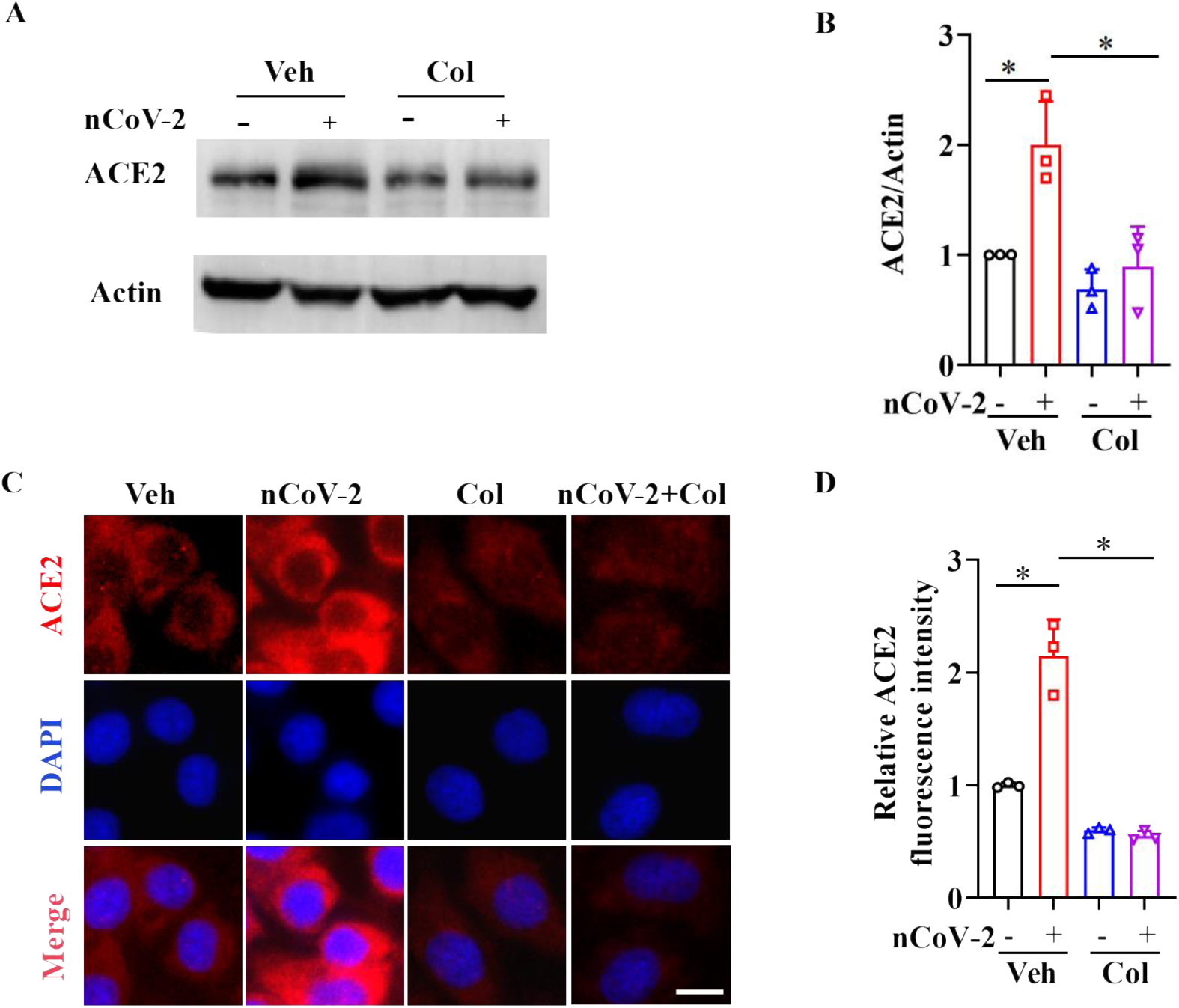
SARS-CoV-2 infection up-regulates ACE2 expression, which is suppressed by colchicine. (**A** and **B**) SARS-CoV-2 infection up-regulated the protein levels of ACE2. Colchicine (20 nM) significantly reduced the protein levels of ACE2 in HPAEpiC cells. The blot is typical of three independent experiments (**A**). Quantification of the ratio of ACE2 to Actin (**B**). These results are means ± SD of three independent experiments. *P < 0.05 (unpaired Student’s t test). (**C**) Representative images of immunofluorescence staining for ACE2. Scale bar: 10 μm. (**D**) Quantification of ACE2 fluorescence intensity. These results are means ± SD of three independent experiments. *P < 0.05 (unpaired Student’s t test). Veh, Vehicle. Col, Colchicine. nCoV-2, SARS-CoV-2. **Figure 1-source data 1** Original uncropped Western blot images in Figure 1A (anti-ACE2 and anti-Actin). **Figure 1-source data 2** PDF containing Figure 1A and original scans of the relevant Western blot analysis (anti-ACE2 and anti-Actin) with highlighted bands and sample labels. **Figure 1-source data 3** Original file for quantification the ratio of ACE2 to Actin in Figure 1B (anti-ACE2 and anti- Actin). **Figure 1-source data 4** Original file for quantification of ACE2 fluorescence intensity in Figure 1D.

As colchicine could inhibit ACE2 expression, we assessed the in vitro inhibitory effect of colchicine on SARS-CoV-2 replication in HPAEpiC cells. After preincubated with colchicine at different concentrations for 1 h, cells were infected with SARS-CoV-2 for 1 h and then cultured in fresh medium for 24 h to measure viral RNA copy numbers by quantitative reverse transcriptase PCR (qPCR). We found that colchicine treatment reduced extracellular SARS-CoV-2 replication in cells (Figure 1-figure supplement 1A). The half-maximal effective concentration (EC_50_) value of colchicine for inhibiting viral replication was 0.2703 μM (Figure 1-figure supplement 1B). These data suggest that inhibition of ACE2 expression by colchicine suppresses SARS-CoV-2 infection.

### SARS-CoV-2 infection regulates the transcription activity of Sp1 and HNF4α

To clarify the molecular mechanism for the regulation of ACE2 expression, we used colchicine, as proof-of-principle. We analyzed a range of signaling pathways in response to SARS-CoV-2 infection in the presence or absence of colchicine using cignal finder 45-pathway reporter array (Manzini et al., 2014, Xu et al., 2021b) (Figure 2A). Of all the signaling pathways tested, there were three transcription factors (Sp1, NF-κB, and GATA) enhanced by SARS-CoV-2 infection, and suppressed by colchicine (Figure 2A). Meanwhile, there were two signaling pathways (HNF4α and estrogen receptor) inactivated by SARS-CoV-2 infection and activated by colchicine treatment (Figure 2A). Thus, these transcription factors are potential candidates for regulating ACE2 expression. We thus analyzed the DNA motifs in 1.5 kb upstream of the transcription start sites (TSS) of ACE2 gene using the MEME program (Bailey et al., 2015). Our data revealed two significantly enriched motifs (Figure 2B), which were annotated as the motif of transcription factors including Sp1 (P = 6.1e-5), HNF4α (P = 4.2e-5). These results implicate that these two transcription factors are likely involved in regulating ACE2 expression. To determine the effects of SARS-CoV-2 and colchicine on the activities of Sp1 and HNF4α, the subcellular distribution of Sp1 and HNF4α were detected using immunofluorescence. Although Sp1 was mainly located in nucleus of HPAEpiC cells (Figure 2-figure supplement 1), only a small portion of total Sp1 was phosphorylated at Thr453 (Figure 2C), which is an indicator of its activation (Milanini-Mongiat et al., 2002). SARS-CoV-2 infection remarkably increased the levels of phospho-Sp1 (Thr453), whereas colchicine treatment significantly inhibited the phosphorylation of Sp1 in the presence or absence of SARS-CoV-2 (Figure 2C and 2D). Unlike Sp1, HNF4α exhibited a dual nuclear and cytoplasmic distribution in HPAEpiC cells under basal conditions (Figure 2E). Whereas SARS-CoV-2 infection induced a nuclear to cytoplasmic shift in the distribution of HNF4α, supplementation with colchicine induced nuclear accumulation of HNF4α in the presence or absence of SARS-CoV-2 (Figure 2E and 2F). These results suggest that SARS-CoV-2 infection results in activation of Sp1 and inactivation of HNF4α, which is reversed by colchicine treatment.

**Figure 2.**
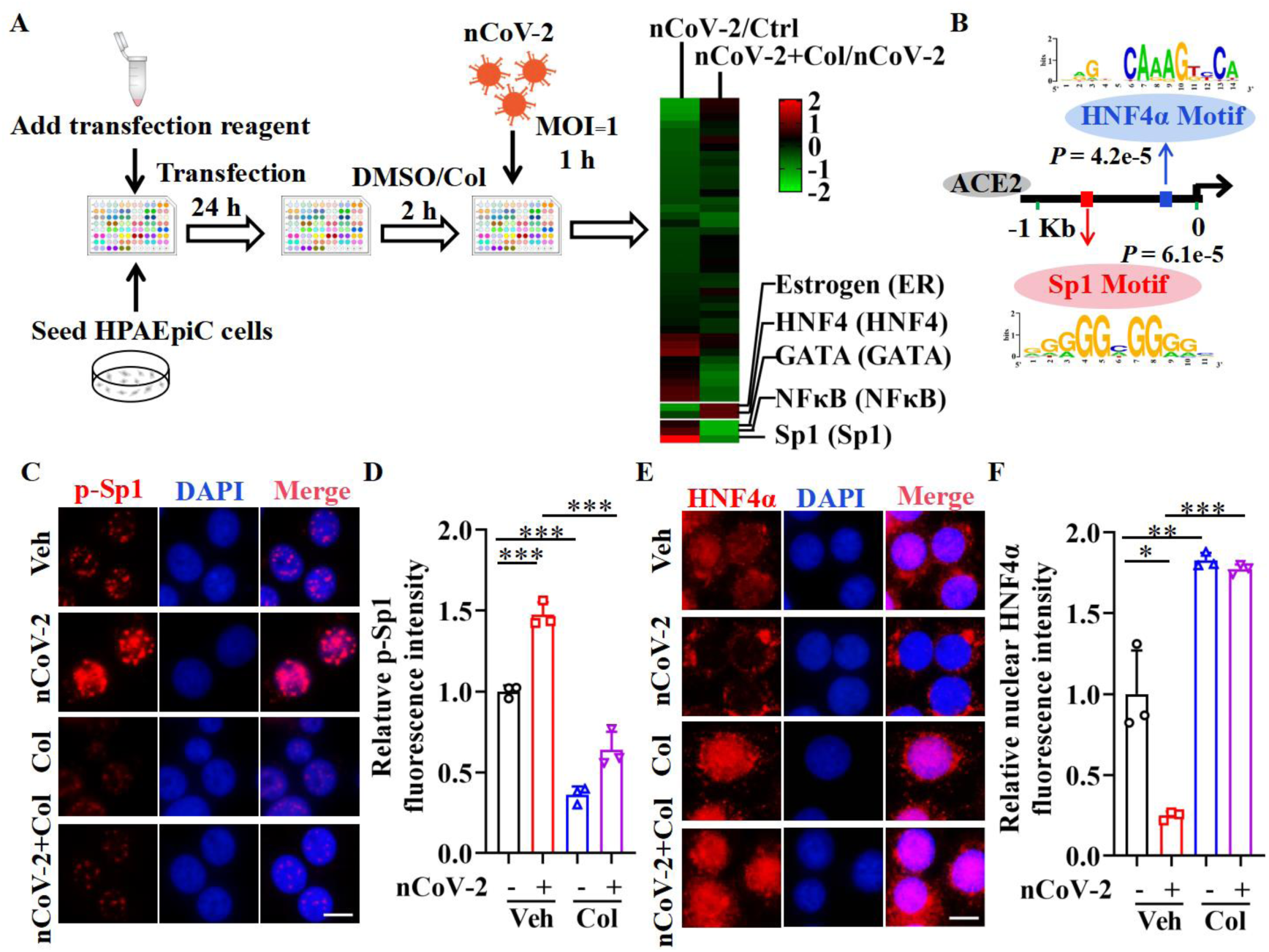
The activation of Sp1 and inactivation of HNF4α are mediated by SARS-CoV-2. (A) Assays for the signaling pathways in response to SARS-CoV-2 infection and colchicine treatment (20 nM) using cignal finder 45-pathway reporter array. (**B**) The Sp1 and HNF4α binding elements in 1.5 kb upstream of the transcription start sites (TSS) of ACE2 gene identified by the MEME program. (**C and D**) SARS-CoV-2 significantly increased the phosphorylation of Sp1 (p-Sp1) in HPAEpiC cells, which was suppressed by treating with colchicine (20 nM). Representative images of immunofluorescence staining for p-Sp1 (**C**). Scale bar: 10 μm. Quantification of p-Sp1 fluorescence intensity (**D**). These results are means ± SD of three independent experiments. \*\*\**P* < 0.001 (unpaired Student’s t test). (**E and F**) SARS-CoV-2 induced cytoplasmic translocation of HNF4α, whereas colchicine (20 nM) promoted its nuclear accumulation in HPAEpiC cells. Representative images of immunofluorescence staining for HNF4α (**E**). Scale bar: 10 μm. Quantification of HNF4α fluorescence intensity (**F**). These results are means ± SD of three independent experiments. \**P* < 0.05, \*\**P* < 0.01, \*\*\**P* < 0.001 (unpaired Student’s t test). Veh, Vehicle. Col, Colchicine. nCoV-2, SARS-CoV-2. **Figure 2-source data 1** Original file for quantification of p-Sp1 fluorescence intensity in Figure 2D. **Figure 2-source data 2** Original file for quantification of HNF4α fluorescence intensity in Figure 2F.

### Sp1 and HNF4α is involved in ACE2 expression

Next, we investigated whether colchicine inhibited ACE2 expression by regulation of Sp1 and HNF4α. First, western blot analysis demonstrated that like colchicine, either treatment with a selective Sp1 inhibitor mithramycin A (MithA) or knockdown of Sp1 by siRNA downregulated the protein levels of ACE2 in HPAEpiC cells (Figure 3A and 3B). However, inhibition of Sp1 by MithA or siSp1 did not further reduce the protein expression of ACE2 downregulated by colchicine. In contrast, both treatment with an HNF4α antagonist BI6015 and knockdown of HNF4α by siRNA upregulated the protein levels of ACE2 (Figure 3C and 3D). Furthermore, colchicine blocked this increase in the protein expression of ACE2 induced by BI6015 or siHNF4α.

**Figure 3.**
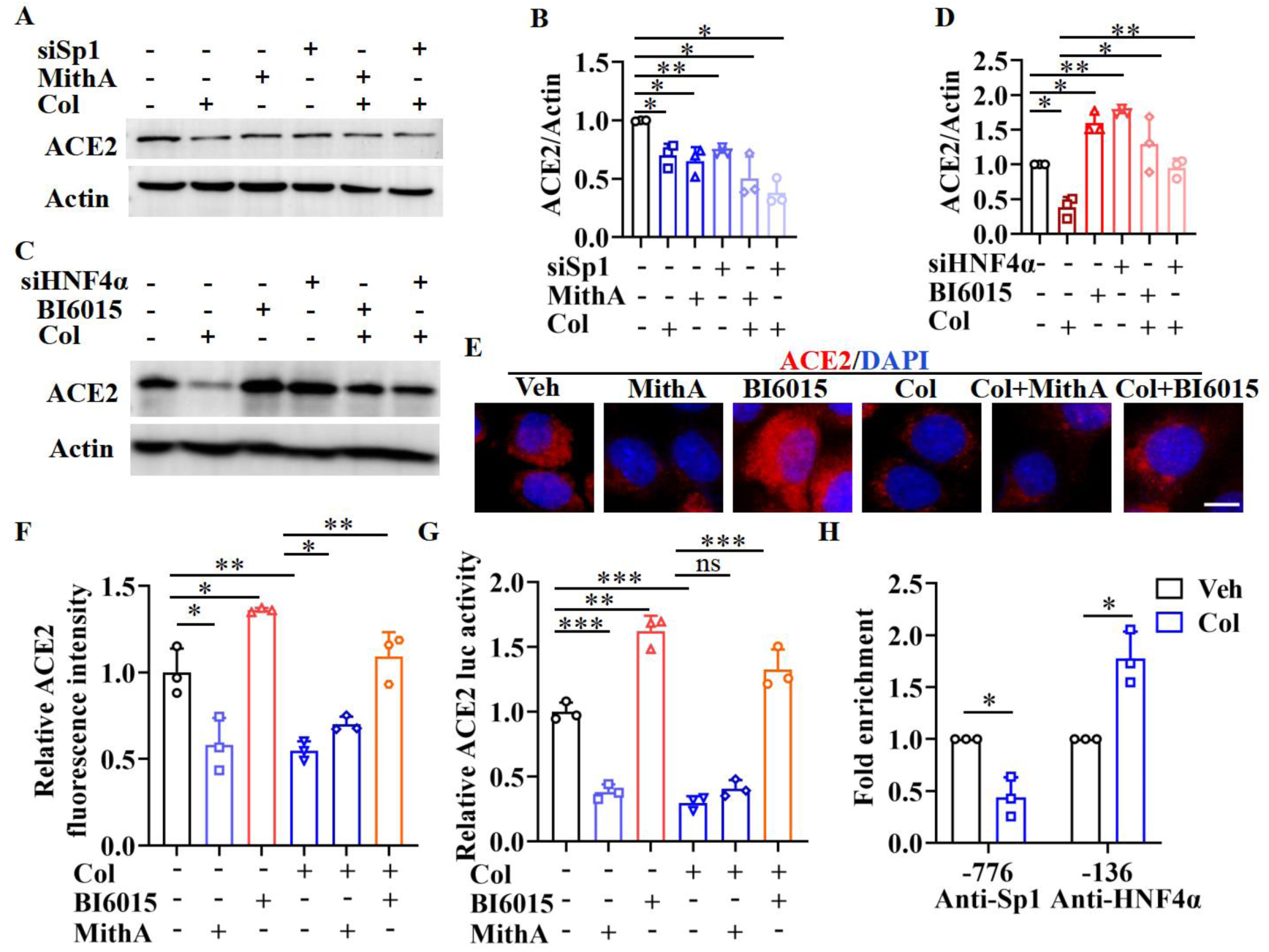
Sp1 and HNF4α act in an opposite way on the regulation of ACE2 expression. (**A and B**) Supplementation with colchicine (20 nM), MithA (100 nM), colchicine + MithA, siSp1, or siSp1 + colchicine significantly suppressed the levels of ACE2 in HPAEpiC cells. The blot is typical of three independent experiments (**A**). Quantification the ratio of ACE2 to Actin (**B**). These results are means ± SD of three independent experiments. \**P* < 0.05, \*\**P* < 0.01 (unpaired Student’s t test). (**C and D**) Supplementation with colchicine (20 nM) significantly suppressed the levels of ACE2 in HPAEpiC cells, which was reversed by treating with BI6015 (20 μM) or siHNF4α. The blot is typical of three independent experiments (**C**). Quantification the ratio of ACE2 to Actin (**D**). These results are means ± SD of three independent experiments. \**P* < 0.05, \*\**P* < 0.01 (unpaired Student’s t test). (**E and F**) Representative images of immunofluorescence staining for ACE2 in HPAEpiC cells (**E**). HPAEpiC cells were treated with MithA (100 nM), BI6015 (20 μM), colchicine (20 nM), colchicine + MithA, or colchicine + BI6015. Scale bar: 10 μm. Quantification of ACE2 fluorescence intensity (**F**). These results are means ± SD of three independent experiments. \**P* < 0.05, \*\**P* < 0.01 (unpaired Student’s t test). (**G**) Luciferase activity analysis of ACE2 promoter in HPAEpiC cells. These results are means ± SD of three independent experiments. \*\**P* < 0.01, \*\*\**P* < 0.001, ns, not significant (unpaired Student’s t test). (**H**) The putative Sp1 and HNF4α binding sites in the promoter regions of ACE2 were detected by ChIP-qPCR with anti-Sp1 and anti-HNF4α antibodies. These results are means ± SD of three independent experiments. \**P* < 0.05. Veh, Vehicle. Col, Colchicine. **Figure 3-source data 1** Original uncropped Western blot images in Figure 3A (anti-ACE2 and anti-Actin). **Figure 3-source data 2** Original uncropped Western blot images in Figure 3C (anti-ACE2 and anti-Actin). **Figure 3-source data 3** PDF containing Figure 3A and 3C and original scans of the relevant Western blot analysis (anti- ACE2 and anti-Actin) with highlighted bands and sample labels. **Figure 3-source data 4** Original file for quantification the ratio of ACE2 to Actin in Figure 3B and 3D (anti-ACE2 and anti-Actin). **Figure 3-source data 5** Original file for quantification of ACE2 fluorescence intensity in Figure 3F. **Figure 3-source data 6** Original file for luciferase activity analysis in Figure 3G. **Figure 3-source data 7** Original file for ChIP analysis in Figure 3H.

Second, immunofluorescence staining of ACE2 revealed that although MithA treatment inhibited the protein expression of ACE2, it did not further reduce the protein expression of ACE2 suppressed by colchicine (Figure 3E and 3F). Supplementation with BI6015 markedly increased the expression of ACE2 in HPAEpiC cells, which was reduced by treated with colchicine (Figure 3E and 3F). Similar results were obtained in A549 cells, a human lung epithelial cell line (Figure 3-figure supplement 1).

Third, using a luciferase reporter gene containing the ACE2 promoter, we observed that supplementation with MithA treatment inhibit the luciferase activity, but did not reduce the luciferase activity in the presence of colchicine (Figure 3G). In contrast, treatment with BI6015 increased the luciferase activity. In addition, chromatin immunoprecipitation (ChIP)–qPCR analysis demonstrated that the binding of Sp1 to the GC box of ACE2 promoter was significantly reduced, whereas the binding of HNF4α to the AGGTCA element was markedly increased after colchicine treatment (Figure 3H). As MithA is capable of inhibiting the expression of ACE2, we tested whether this compound could inhibit SARS-CoV-2 infection in vitro. Our results demonstrated that MithA inhibited SARS-CoV-2 replication with EC50 of 0.1948 μM in HPAEpiC cells (Figure 3-figure supplement 2). Taken together, these results suggest that Sp1 and HNF4α act in an opposite way on the regulation of ACE2 expression at the transcription level to affect the susceptibility of cells to SARS-CoV-2 infection.

### Sp1 and HNF4α antagonized each other

Interestingly, we found that treatment with the Sp1 inhibitor MithA significantly suppressed the phosphorylation of Sp1, whereas treatment with the HNF4α antagonist BI6015 led to an increase in the levels of phospho-Sp1 (Thr453) (Figure 4A and 4B). Conversely, BI6015 treatment induced cytoplasmic translocation of HNF4α, whereas MithA treatment promoted nuclear accumulation of HNF4α (Figure 4C and 4D). Likewise, western blotting analysis revealed that the phosphorylation level of Sp1 was increased after knockdown of HNF4α by siRNA (Figure 4E and 4F). By contrast, knockdown of Sp1 did not affect the levels of total HNF4α. These data led us to test whether Sp1 could interact with HNF4α. Co- immunoprecipitation assay (co-IP) confirmed the interaction between Sp1 and HNF4α in HPAEpiC cells (Figure 4G). These data indicate that Sp1 and HNF4α antagonize each other probably by protein-protein interaction.

**Figure 4.**
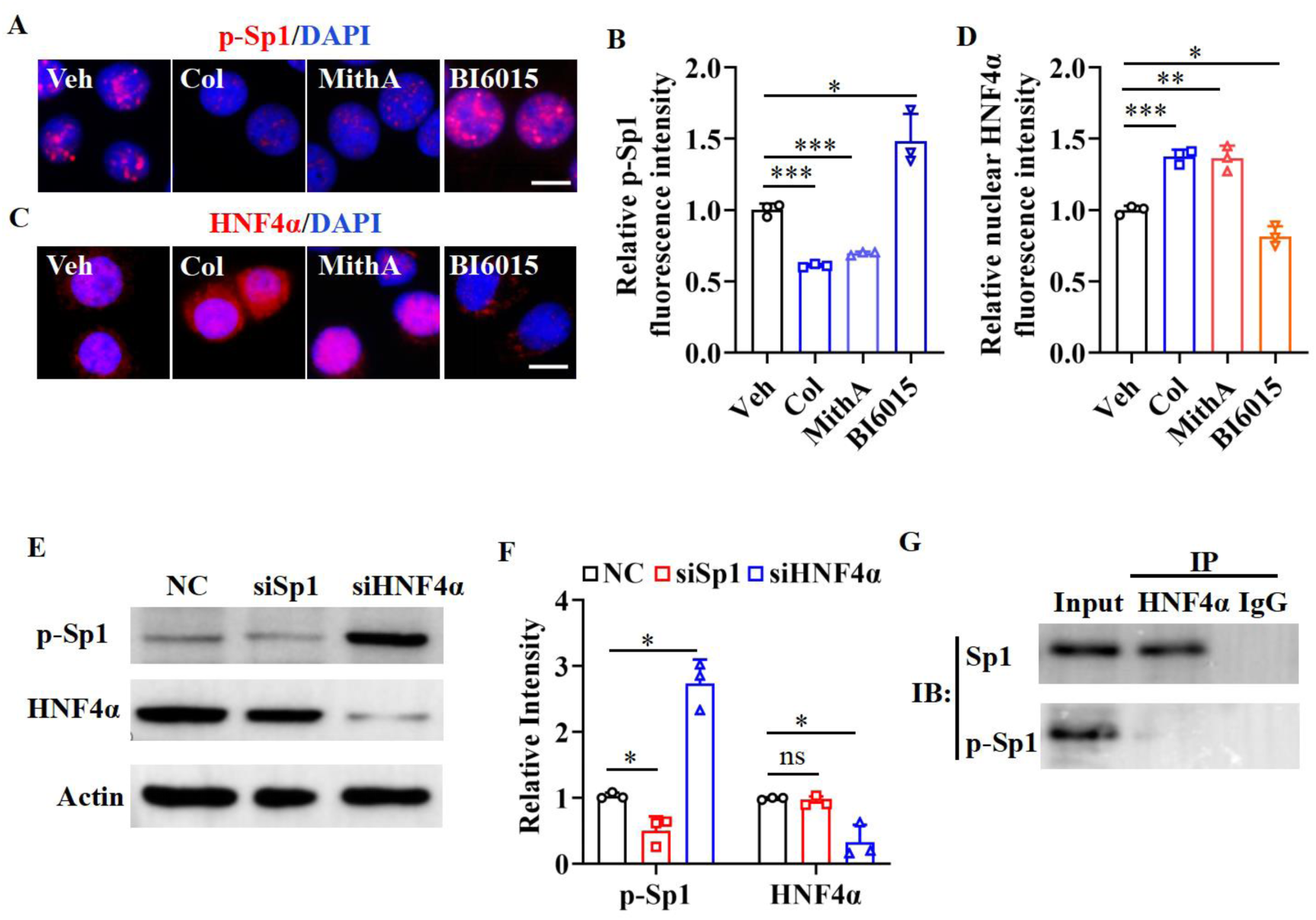
Sp1 and HNF4α antagonize each other via protein-protein interaction. **(A and B)** Supplementation with colchicine (20 nM) and MithA (100 nM) significantly suppressed the phosphorylation of Sp1 (p-Sp1) in HPAEpiC cells, which was reversed by treating with BI6015 (20 μM). Representative images of immunofluorescence staining for p-Sp1 (**A**). Scale bar: 10 μm. Quantification of p-Sp1 fluorescence intensity (**B**). These results are means ± SD of three independent experiments. \**P* < 0.05, \*\*\**P* < 0.001 (unpaired Student’s t test). (**C and D**) Supplementation with colchicine (20 nM) and MithA (100 nM) promoted nuclear accumulation of HNF4α in HPAEpiC cells, which was inhibited by treating with BI6015 (20 μM). Representative images of immunofluorescence staining for HNF4α (**C**). Scale bar: 10 μm. Quantification of HNF4α fluorescence intensity (**D**). These results are means ± SD of three independent experiments. \**P* < 0.05, \*\**P* < 0.01, \*\*\**P* < 0.001 (unpaired Student’s t test). (**E and F**) The phosphorylation levels of Sp1 and the total protein levels of HNF4α were measured in HPAEpiC cells by Western blotting. The blot is typical of three independent experiments (**E**). Quantification of the ratio of p-Sp1 or HNF4α to Actin (**F**). These results are means ± SD of three independent experiments. \**P* < 0.05, ns, not significant (unpaired Student’s t test). (**G**) The interaction between Sp1 and HNF4α measured by Co-immunoprecipitation assay (co-IP) in HPAEpiC cells. NC, Negative control. Veh, Vehicle. Col, Colchicine. **Figure 4-source data 1** Original file for quantification of p-Sp1 fluorescence intensity in Figure 4B. **Figure 4-source data 2** Original file for quantification of HNF4α fluorescence intensity in Figure 4D. **Figure 4-source data 3** Original uncropped Western blot images in Figure 4E (anti-p-Sp1, anti-HNF4α and anti-Actin). **Figure 4-source data 4** PDF containing Figure 4E and 4G and original scans of the relevant Western blot analysis (anti-p-Sp1, anti-HNF4α and anti-Actin) with highlighted bands and sample labels. **Figure 4-source data 5** Original file for quantification the ratio of p-Sp1 and HNF4α to Actin in Figure 4F (anti-p-Sp1, anti-HNF4α and anti-Actin). **Figure 4-source data 6** Original uncropped Western blot images in Figure 4G.

### Colchicine reduces ACE2 expression by inhibiting the PI3K/AKT signaling pathway

It has been reported that the PI3K/AKT signaling pathway is activated by SARS-CoV-2 infection (Callahan et al., 2021, Sun et al., 2021, Klann et al., 2020). Consistent with these observations, we found that SARS-CoV-2 infection enhanced phosphorylation of AKT at Ser473 and Thr308, which is required for its activation (Su et al., 2011) (Figure 5A-5C). However, colchicine treatment substantially inhibited SARS-CoV-2-induced AKT phosphorylation (Figure 5A-5C). Furthermore, either knockdown of AKT by siRNA or treatment with two PI3K/AKT inhibitors (LY294002 and wortmannin) downregulated the expression of ACE2 in HPAEpiC cells (Figure 5D-5G). Finally, we found that LY294002 and wortmannin inhibited SARS-CoV-2 infection in HPAEpiC cells, with EC_50_ of 0.2381 μM and 0.04228 μM, respectively (Figure 5-figure supplement 1). These data suggest that the PI3K/AKT signaling pathway is required for SARS-CoV-2 infection.

**Figure 5.**
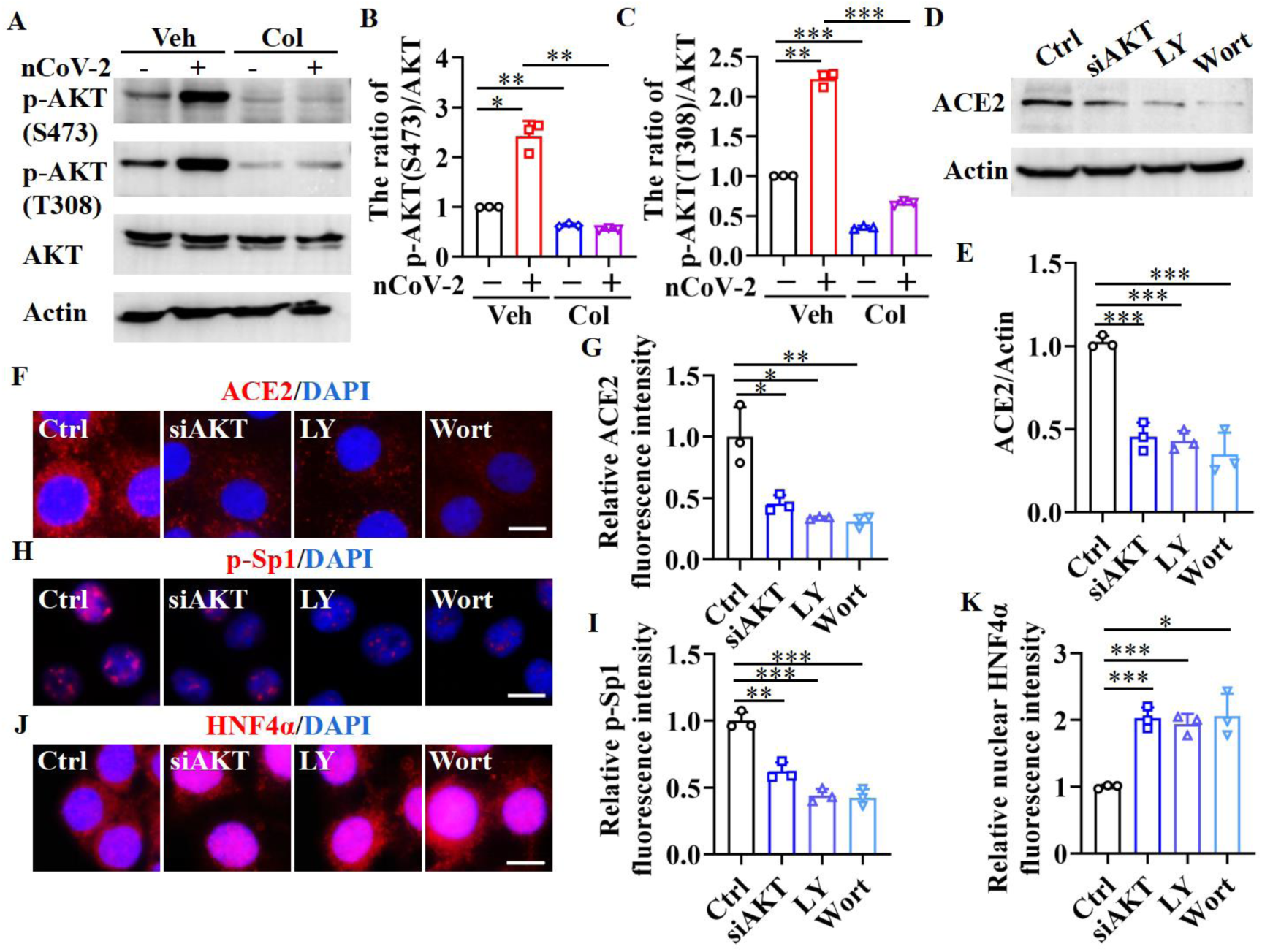
Sp1 and HNF4α are the downstream of PI3K/AKT signaling pathway. (**A-C**) The phosphorylation levels of AKT (S473 and T308) were measured in HPAEpiC cells by Western blotting. The blot is typical of three independent experiments (**A**). Quantification of the ratio of p-AKT (S473) (**B**) or p-AKT (T308) (**C**) to Actin. These results are means ± SD of three independent experiments. \**P* < 0.05, \*\**P* < 0.01, \*\*\**P* < 0.001 (unpaired Student’s t test). (**D-G**) Supplementation with LY294002 (30 μM) and wortmannin (50 nM), or siAKT significantly suppressed the levels of ACE2 in HPAEpiC cells. The blot is typical of three independent experiments (**D**). Quantification of the ratio of ACE2 to Actin (**E**). These results are means ± SD of three independent experiments. \*\*\**P* < 0.001 (unpaired Student’s t test). Representative images of immunofluorescence staining for ACE2 (**F**). Scale bar: 10 μm. Quantification of ACE2 fluorescence intensity (**G**). These results are means ± SD of three independent experiments. \**P* < 0.05, \*\**P* < 0.01 (unpaired Student’s t test). (**H and I**) Supplementation with LY294002 (30 μM) and wortmannin (50 nM), or siAKT significantly suppressed the phosphorylation of Sp1 (p-Sp1) in HPAEpiC cells. Representative images of immunofluorescence staining for p-Sp1 (**H**). Scale bar: 10 μm. Quantification of p-Sp1 fluorescence intensity (**I**). These results are means ± SD of three independent experiments. \*\**P* < 0.01, \*\*\**P* < 0.001 (unpaired Student’s t test). (**J and K**) Supplementation with LY294002 (30 μM) and wortmannin (50 nM), or siAKT promoted nuclear accumulation HNF4α in HPAEpiC cells. Representative images of immunofluorescence staining for HNF4α (**J**). Scale bar: 10 μm. Quantification of HNF4α fluorescence intensity (**K**). These results are means ± SD of three independent experiments. \**P* < 0.05, \*\*\**P* < 0.001 (unpaired Student’s t test). Ctrl, Control. Veh, Vehicle. Col, Colchicine. LY, LY294002. Wort, wortmannin. nCoV-2, SARS- CoV-2. **Figure 5-source data 1** Original uncropped Western blot images in Figure 5A (anti-p-AKT (S473), anti-p-AKT (T308) and anti-Actin). **Figure 5-source data 2** PDF containing Figure 5A and 5D and original scans of the relevant Western blot analysis with highlighted bands and sample labels. **Figure 5-source data 3** Original file for quantification the ratio of p-AKT to AKT in Figure 5B and 5C. **Figure 5-source data 4** Original uncropped Western blot images in Figure 5D (anti-ACE2 and anti-Actin). **Figure 5-source data 5** Original file for quantification the ratio of ACE2 to Actin in Figure 5E. **Figure 5-source data 6** Original files for quantification of ACE2, p-Sp1 and HNF4α fluorescence intensity in Figure 5G, 5I and 5K.

The PI3K/AKT signaling pathway plays an important role in regulating the transcriptional activity of Sp1 and HNF4α (Adapala et al., 2019, Gomez-Villafuertes et al., 2015, Zhao et al., 2015, Li et al., 2019). Activation of AKT promotes the stability and localization of Sp1 by phosphorylating Sp1 at threonine sites 453 and 739 (Adapala et al., 2019, Gomez-Villafuertes et al., 2015, Zhao et al., 2015), and prevents the nuclear translocation of HNF4α (Li et al., 2019). Consistent with these observations, we found that suppression of AKT by siRNA or its inhibitors significantly inhibited the accumulation of phospho-Sp1 in the nucleus, but remarkably promoted nuclear translocation of HNF4α (Figure 5H-5K). Taken together, these data indicate that inhibition of the PI3K/AKT signaling pathway is required for downregulation of ACE2 expression mediated by colchicine via regulation of Sp1 and HNF4α transcription activities.

### Inhibition of Sp1 reduces the viral load and damage to the respiratory and renal systems

Finally, we tested whether inhibition of Sp1 by colchicine and MithA could inhibit the replication of SARS-CoV-2 in vivo by using Syrian hamsters (*Mesocricetus auratus*), an animal model used for the study of COVID-19 pneumonia and the evaluation of therapeutics (Chan et al., 2020, Choudhary et al., 2022, Muñoz-Fontela et al., 2020, Rosenke et al., 2021, Sia et al., 2020). Two groups of hamsters were intranasally infected with SARS-CoV-2 at a dose of 10^4^ TCID_50_ (Tissue culture infective dose). One hour later, hamsters were inoculated intraperitoneally with colchicine or MithA at 0.2 mg/kg, respectively. A mock group was treated with vehicle using the same route and timing (Figure 6-figure supplement 1). These animals were dosed every 24 h with either colchicine or MithA at 0.2 mg/kg, respectively. Lung and trachea samples were collected at 3 days post-infection for assessing viral RNA and ACE2 expression. Immunofluorescence analysis demonstrated that the expression of ACE2 was significantly upregulated in the lung and trachea of hamsters infected with SARS-CoV-2, compared to that of animals without infection (control group) (Figure 6A and 6B; Figure 6-figure supplement 2A and Figure 6-figure supplement 2B). However, treatment with colchicine or MithA substantially inhibited the expression of ACE2 in both of the lung and trachea of hamsters infected with SARS-CoV-2. Based on immunofluorescence analysis for SARS-CoV- 2 nucleocapsid and qPCR, we found that treatment with colchicine or MithA reduced viral replication in both of the lung and trachea of hamsters (Figure 6C-6F; Figure Figure 6-figure supplement 2C and Figure 6-figure supplement 2D), respectively.

**Figure 6.**
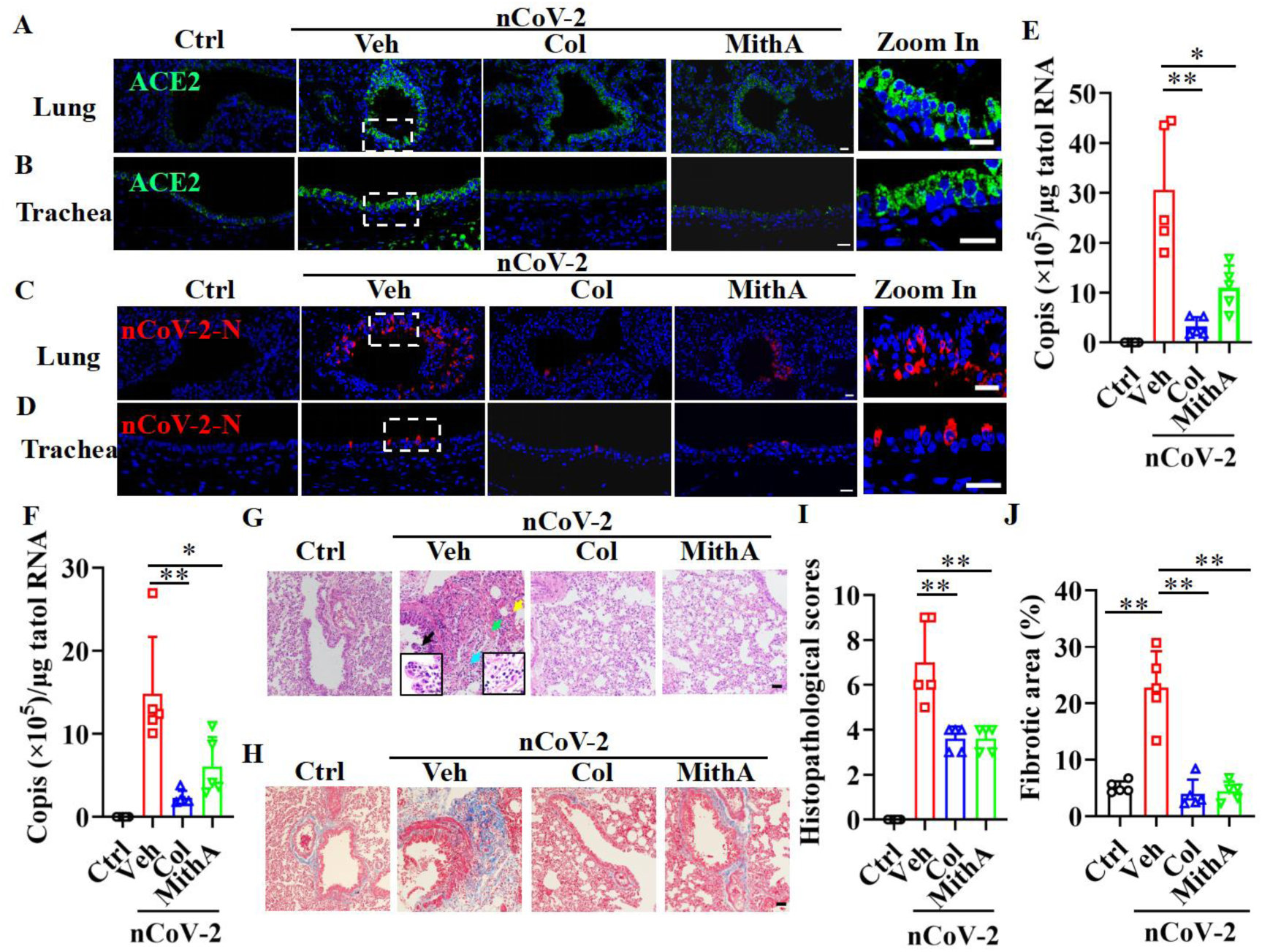
Inhibition of Sp1 inhibits the replication of SARS-CoV-2 and reduces lung pathology in Syrian hamsters. (**A and B**) Treatment of either colchicine or MithA inhibited the expression of ACE2 in the lung and trachea of hamsters infected with SARS-CoV-2. Representative images of immunofluorescence staining of ACE2 in the lung (**A**) and trachea (B) of hamsters. The parts on the right side are high-power images. Scale bar: 20 μm. (**C-F**) Treatment of either colchicine or MithA inhibited the replication of SARS-CoV-2 in the lung and trachea of hamsters. Representative images of immunofluorescence staining of SARS- CoV-2-N in the lung (**C**) and trachea (**D**) of hamsters. The parts on the right side are high- power images. Scale bar: 20 μm. Viral loads of SARS-CoV-2 were measured in the lung (**E**) and trachea (**F**) of hamsters by qPCR (n = 5 each group). Error bars show means ± SD. \**P* < 0.05, \*\**P* < 0.01 (unpaired Student’s t test). (**G-J**) Supplementation with either colchicine or MithA attenuated histopathological damage in the lung of hamsters infected with SARS-CoV-2. Representative images of haematoxylin and eosin (HE) staining in lung of hamsters infected with SARS-CoV-2 at 3 dpi (**G**). Bronchial epithelial cell necrosis and pyknosis (black arrow), edema, loose arrangement of muscle fibers, and massive lymphocyte infiltration (blue arrow), extensive hemorrhage (green arrow), and alveolar wall thickening (yellow arrow). The parts on the lower side are high-power images of black arrow and blue arrow, respectively. Scale bar: 40 μm. Representative images of masson staining in lung of hamsters infected with SARS-CoV-2 at 3 dpi (**H**). Scale bar: 40 μm. Summary of lung lesion scoring in different groups at 3 dpi (n=5 each group) (**I**). Error bars show means ± SD. \**P* < 0.05 (unpaired Student’s t test). Quantitative analysis of fibrotic area in lung tissues (**J**). Error bars show means ± SD. \*\**P* < 0.01 (unpaired Student’s t test). Ctrl, Control. Veh, Vehicle. Col, Colchicine. nCoV-2, SARS-CoV-2. **Figure 6-source data 1** Original file for determination of viral load in Figure 6E. **Figure 6-source data 2** Original file for determination of viral load in Figure 6F. **Figure 6-source data 3** Original file for lesion scores in Figure 6I. **Figure 6-source data 4** Original file for quantitative analysis of fibrotic area in Figure 6J.

Next, histopathological analysis of the lung tissue on day 3 post-infection showed pulmonary lesions consisting of necrosis and nuclear pyknosis in bronchial epithelial cells, massive hemorrhage in the alveolar space, a significant invasion of inflammatory cells, and edema (Figure 6G and 6I). In addition, alveolar walls were significantly thickened. While the treatment groups had similar lesions, they were much less severe than the mock group. Using Masson’s trichrome staining, we observed collagen deposition, a hallmark of fibrosis, in the lung of infected hamsters (Figure 6H and 6J). However, treatment with colchicine or MithA effectively reversed lung fibrosis. Taken together, these results suggest that colchicine and MithA antagonize SARS-CoV-2 replication in the lung and trachea, and attenuate histopathological damage in the lung.

Although severe SARS-CoV-2-associated acute kidney injury serves as an independent risk factor for in-hospital death in patients (Nadim et al., 2020), whether SARS-CoV-2 can directly infect the kidney remains unclear (Wysocki et al., 2021, Smith and Akilesh, 2021). The prevailing evidence seems to favor indirect means of kidney injury in SARS-CoV-2 (Smith and Akilesh, 2021). Based on immunofluorescence analysis, the presence of massive SARS-CoV- 2 was observed in the kidney of infected hamsters, which was slightly reduced by treatment with colchicine or MithA (Figure 7A and 7B). Immunofluorescence analysis showed that treatment with the two drugs could effectively reduce the expression of ACE2 (Figure 7C and 7D). Histopathological analysis indicated a significant damage in the kidney of hamsters infected with SARS-CoV-2, including renal tubular epithelial cell nuclear pyknosis, brush border disappearance; renal interstitial vascular congestion, inflammatory cell infiltration; glomerular atrophy (Figure 7E and 7F). By contrast, the glomeruli were evenly distributed, and the structure was intact; the tubular epithelial cells were normal, and the brush borders were neatly arranged in these drug-treated animals. There was no obvious abnormality in the medulla, and no obvious hyperplasia of the renal interstitium. Taken together, treatment with colchicine and MithA leads to a significant improvement in renal histology.

**Figure 7.**
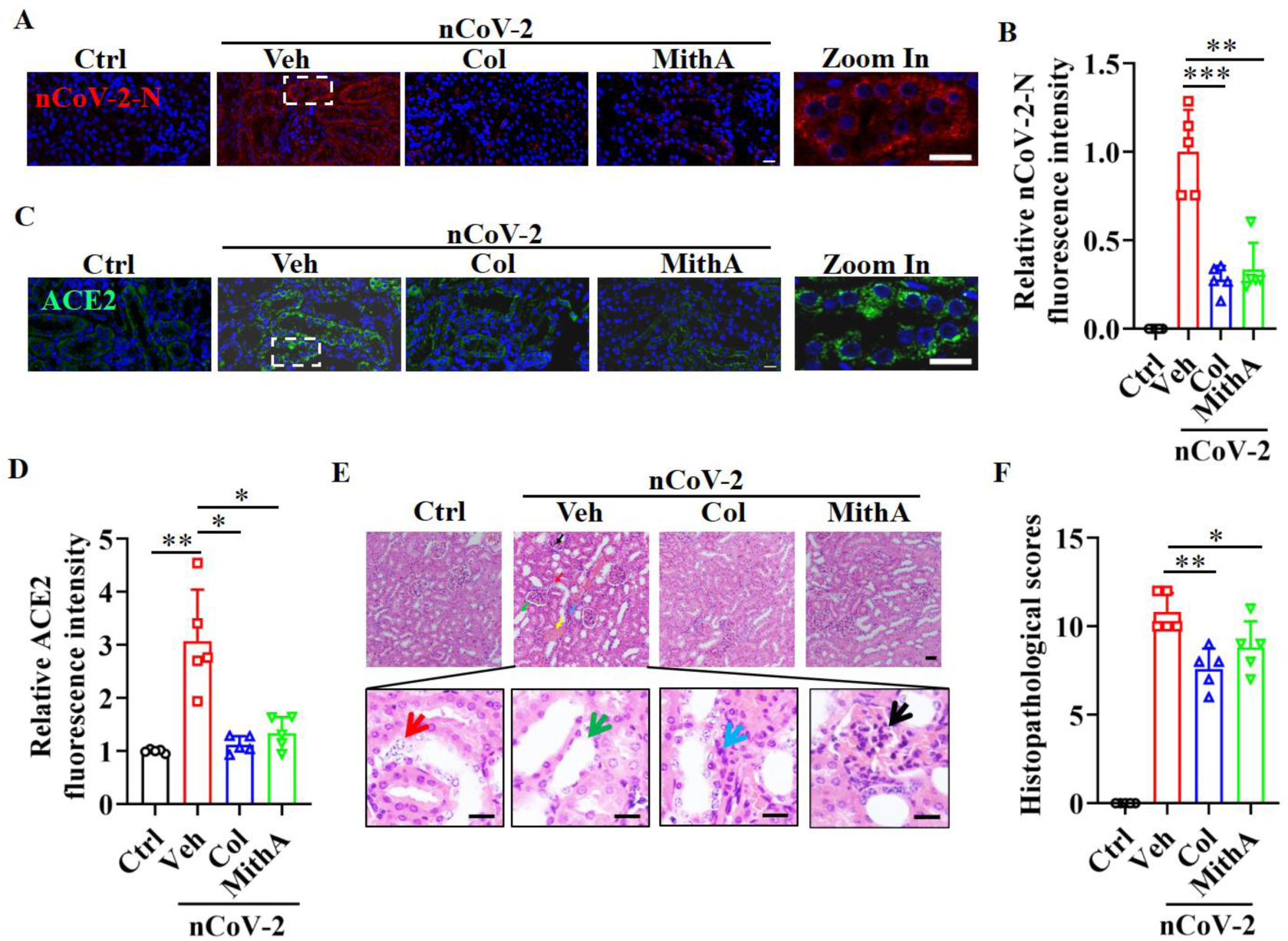
Inhibition of Sp1 inhibits the replication of SARS-CoV-2 and reduces kidney pathology in Syrian hamsters. (**A and B**) Treatment of either colchicine or MithA inhibited the replication of SARS-CoV-2 in the kidney of hamsters. Representative images of immunofluorescence staining of SARS-CoV-2-N in the kidney of hamsters (**A**). The parts on the right side are high-power images. Scale bar: 20 μm. Quantification of SARS-CoV-2-N fluorescence intensity in the kidney of hamsters (**B**). Error bars show means ± SD. \*\**P* < 0.01, \*\*\**P* < 0.001 (Unpaired Student’s t test). (**C and D**) Treatment of either colchicine or MithA inhibited the expression of ACE2 in the kidney of hamsters infected with SARS-CoV-2. Representative images of immunofluorescence staining of ACE2 in the kidney of hamsters (**C**). The parts on the right side are high-power images. Scale bar: 20 μm. Quantification of ACE2 fluorescence intensity in the kidney of hamsters (**D**). Error bars show means ± SD. \**P* < 0.05, \*\**P* < 0.01 (Unpaired Student’s t test). (**E and F**) Supplementation with either colchicine or MithA attenuated histopathological damage in the kidney of hamsters infected with SARS- CoV-2. Representative images of haematoxylin and eosin (HE) staining in the kidney of hamsters infected with SARS-CoV-2 at 3 dpi (**E**). Histopathology of kidney showed renal interstitial vascular congestion (black arrow), renal tubular epithelial cell nuclear pyknosis (red arrow), brush border disappearance (green arrow), renal interstitial inflammatory cell infiltration (blue arrow), and lomerular atrophy (yellow arrow). The parts on the lower side are high-power images of red arrow, green arrow, and blue arrow, respectively. Scale bar: 40 μm. Summary of kidney lesion scoring in different groups at 3 dpi (n=5 each group) (**F**). Error bars show means ± SD. \**P* < 0.05, \*\**P* < 0.01 (Unpaired Student’s t test). Ctrl, Control. Veh, Vehicle. Col, Colchicine. nCoV-2, SARS-CoV-2. **Figure 7-source data 1** Original file for quantification of SARS-CoV-2-N fluorescence intensity in Figure 7B. **Figure 7-source data 2** Original file for quantification of ACE2 fluorescence intensity in Figure 7D. **Figure 7-source data 3** Original file for lesion scores in Figure 7F.

## Discussion

Based on our study and our current understanding of SARS-CoV-2 infection, we propose a model for how SARS-CoV-2 infection induces ACE2 expression through a mechanism involving two transcription factors, Sp1 and HNF4α, in host cells. Under normal conditions, Sp1 is mainly located in nucleus, whereas HNF4α exhibit a dual nuclear and cytoplasmic distribution in cells. The two transcription factors regulate ACE2 expression in an opposite manner, leading to a basal expression of ACE2. After SARS-CoV-2 infection, the PI3K/AKT signaling pathway is activated (Klann et al., 2020). This signaling pathway in turn promotes the transcription activity of Sp1 by increasing its phosphorylation in nucleus, and suppresses the transcription activity of HNF4α by inducing its cytoplasmic translocation. A disruption of this balance leads to the upregulation of ACE2. Since ACE2 is the receptor for the SARS-CoV- 2 viral entry, induction of ACE2 expression may represent an effective tactic employed by the virus to facilitate its own propagation.

Multi-omics approaches have recently been applied to understand host responses to SARS-CoV-2, thus accelerating drug development and drug repositioning against COVID-19 (Kamel et al., 2021, Lu et al., 2022, Chu et al., 2021, Ho et al., 2021, Klann et al., 2020). For instance, phosphoproteomic analysis of SARS-CoV-2-infected cells has revealed the activation of growth factor receptor (GFR) and its downstream pathways including the RAF/MEK/ERK MAPK signaling and PI3K/AKT/mTOR signaling in SARS-CoV-2-infected cells (Klann et al., 2020). Inhibition of the GFR signaling by using prominent anti-cancer drugs, such assorafenib (the RAF inhibitor), RO5126766 (the dual RAF/MEK inhibitor), and pictilisib (the PI3K inhibitor), and omipalisib (the dual PI3K and mTOR inhibitor), prevents SARS-CoV-2 replication in cells. By screening a panel of 45 signaling pathways in SARS-CoV-2-infected HPAEpiC cells in the presence or absence of colchicine, we showed that two transcription factors Sp1 and HNF4α are involved in regulation of ACE2 expression in an opposite manner. Sp1 positively and HNF4α negatively regulate the expression of ACE2 at the transcriptional levels. The PI3K/AKT signaling pathway, which is activated by SARS-CoV-2 infection (Klann et al., 2020), is a crucial node for induction of ACE2 expression by promoting the transcriptional activity of Sp1 and reducing the transcriptional activity of HNF4α.

Our data indicate that down-regulation of ACE2 expression by inhibiting Sp1 activity attenuates SARS-CoV-2 replication in the lung and trachea of Syrian hamsters. Importantly, this inhibition of viral replication in these tissues is associated with markedly reduced lung pathology. Acute kidney injury is another extrapulmonary manifestation of severe COVID-19 (Legrand et al., 2021). However, a direct causal association between SARS-CoV-2 infection and the development of acute kidney injury has been controversial (Legrand et al., 2021, Smith and Akilesh, 2021). Recently, Jansen et al. have demonstrated that SARS-CoV-2 directly infects kidney cells by using human-induced pluripotent stem cell-derived kidney organoids (Jansen et al., 2022). Our data show that colchicine treatment effectively suppresses the expression of ACE2 in the proximal tubule of kidney, which is accompanied by reduced SARS-CoV-2- induced kidney injury in Syrian hamsters. Our results support the notion that ACE2 is a promising therapeutic target in the fight against COVID-19 (Monteil et al., 2020). For example, proxalutamide, a potent androgen receptor antagonist, can suppress the expression of ACE2, which is induced by androgens through androgen receptor (Qiao et al., 2021). Moreover, inhibition of RIPK1 reduces viral load in cultured human lung organoids infected by SARS-CoV-2, which is accompanied by downregulating the transcriptional induction of ACE2 (Xu et al., 2021a). Very recently, Brevini et al. reveal that FXR is involved in the regulation of ACE2 expression in multiple tissues. Inhibition of FXR activity by UDCA downregulates ACE2 expression and reduces SARS-CoV-2 infection in vitro, in vivo and ex vivo (Brevini et al., 2022).

While studying the effect of Sp1 and HNF4α inhibitors on ACE2 expression, we unexpectedly observed that inhibition of Sp1 by its inhibitor MithA induces nuclear accumulation of HNF4α, whereas inhibition of HNF4α by its antagonist BI6015 increases the phosphorylation levels of Sp1 in nucleus. How do the two transcription factors antagonize each other? Although the action of MithA is to interfere with Sp1 binding to its consensus site (Lee et al., 2011), the Sp1 inhibitor is capable of inducing proteasome-dependent Sp1 degradation (Lee et al., 2012, Choi et al., 2014). This may explain why MithA treatment significantly suppresses the phosphorylation levels of Sp1 in our experiments. Likewise, BI6015 can suppress HNF4α expression, although the function of the HNF4α antagonist is to repress HNF4α DNA binding (Kiselyuk et al., 2012). This again provides a ready explanation of our observation that BI6015 could inhibit nuclear accumulation of HNF4α induced by colchicine. More importantly, our results demonstrate that non-phosphorylated Sp1, but not phosphorylated Sp1, can interact with HNF4α. On the basis of these results, one may speculate that the interaction between HNF4α and Sp1 may block Sp1 phosphorylation. Reduced protein levels of HNF4α by BI6015 leads to release of more Sp1, which is ready for phosphorylation by AKT. Phosphorylated Sp1 then binds to the GC box and upregulates ACE2 expression. On the other hand, a decrease in Sp1 protein levels by MithA leads to release of more HNF4α. This transcription factor in turn inhibits ACE2 expression by binding to the HNF4α-specific binding motif. These data may help us to further understand how Sp1 and HNF4α antagonize each other’s transcription activity. Clearly, the underlying mechanisms need to be further investigated in light of our current results.

Increased expression of ACE2 in the airway and lung contributes to the severity of COVID-19 symptoms in elderly patients (Inde et al., 2021). At present, the newly emerging SARS-CoV-2 Omicron variants have become the dominant strain worldwide. The Omicron spike protein has six-fold to nine-fold increased affinity for binding to ACE2 (Yin et al., 2022). Various types of available vaccines, which are based on the original strain of SARS-CoV-2, are ineffective against Omicron variants (Wilhelm et al., 2022, Kuhlmann et al., 2022). Our findings may contribute to knowledge for developing a new host-targeting approach for the fight against COVID-19, especially for the elderly patients with SARS-CoV-2 Omicron variant infection.

## Material and Methods

### Cell culture

Immortalized human alveolar epithelial cells (HPAEpiCs) were generated from human lung tissue type II pneumocytes (purchased from the ScienCell Research Laboratory (San Diego, CA)) and maintained in RPMI 1640 medium (01-100-1ACS, Biological Industries, Israel) supplemented with 10 % fetal bovine serum (FBS) (C04001-050X10, VivaCell, Shanghai, China), and 1 % penicillin-streptomycin. A549 cells were maintained in DMEM/F12 medium (C3130-0500, VivaCell) containing 10 % FBS, and 1 % penicillin-streptomycin. All cells were cultured in 5 % CO2, 95 % air incubator at 37 °C.

HPAEpiCs and A549 cells were treated with colchicine (C804812, Macklin, Shanghai, China), mithramycin A (MithA, A600668, Sangon Biotech, Shanghai, China), BI6015 (HY- 108469, Med Chem Express, Shanghai, China), LY294002 (HY-10108, Med Chem Express), or wortmannin (HY-10197, Med Chem Express). DMSO was used as a control. Cells were infected with SARS-CoV-2 at MOI of 1 for 1 h.

### SARS-CoV-2

The SARS-CoV-2 strain (accession number: NMDCN0000HUI) was provided by the Guangdong Provincial Center for Disease Control and Prevention (Guangzhou, China). The virus was propagated in African green monkey kidney epithelial cells (Vero-E6) (ATCC, No. 1586) and titrated. All the infection experiments were performed in a biosafety level-3 (BLS-3) laboratory.

### Half-maximal effect concentration (EC_50_)

HPAEpiC cells were seeded at a density of 1.6×10^4^ cell/well in 48-well plates and grown overnight. Cells were then infected with SARS-CoV-2 at MOI of 1. At the same time, the test compounds were added to the wells with different concentrations. After 1 h of incubation at 37 °C, the virus-drug mixture was removed and washed 3 times to remove free virus with PBS, replaced with fresh medium containing compounds. After 48 h, the supernatants were collected to extract viral RNA for RT-qPCR analysis. The EC_50_ values were calculated by using a dose-response model in GraphPad Prism 8.0 software (GraphPad Software Inc., La Jolla, CA).

### Cellular antiviral activity assay

HPAEpiC cells were seeded at a density of 4×10^5^ cell/well in 24-well plates and grown overnight. After preincubated with the test compounds for 2 h, the cells were infected with SARS-CoV-2 at an MOI of 1. After 1 h of incubation at 37 °C, the virus-drug mixtures were replaced with fresh medium containing compounds. In 24 h, cells were collected to extract total RNA and total cell protein. Viral RNA was quantified by THUNDERBIRD Probe One-step RT-qPCR Kit (QRZ-101, Toyobo, Shanghai, China). TaqMan primers for SARS-CoV-2 are 5’- GGG GAA CTT CTC CTG CTA GAA T-3’ and 5’-CAG ACA TTT TGC TCT CAA GCT G-3’ with SARS-CoV-2 probe FAM-TTG CTG CTG CTT GAC AGA TT-TAMRA-3’.

### Quantitative real-time PCR

The total RNA was extracted from Trachea and lung tissues using the RNAiso Plus (Takara, Dalian, China). Total RNA was extracted from cells with TRIzol™ Reagent (R1100, Solarbio, Shanghai, China), and reverse-transcribed into cDNA using FastKing RT Kit (KR116, TIANGEN, Beijing, China). qPCR analysis was performed using SuperReal PreMix Plus (SYBR Green) (FP205, TIANGEN) on a Roche LightCycler 480 System (Roche Applied Science, Mannheim, Germany). Primers used for qPCR were followed: ACE2 (Forward 5’- GGG ATC AGA GAT CGG AAG AAG AAA-3’; Reverse 5’-AGG AGG TCT GAA CAT CAT CAG TG-3’); ACTB (Forward 5’-CCC TGG ACT TCG AGC AAG AG-3’; Reverse 5’-ACT CCA TGC CCA GGA AGG AA-3’). The relative mRNA expression levels of ACE2 were assessed by the 2^-ΔΔCt^ method. ACTB was used to calculate relative expression normalized to an internal control.

### 45-Pathway Reporter Array

Cignal Finder 45-Pathway Reporter Arrays (CCA-901, Qiagen, Hilden, Germany) were used according to the manufacturer’s instructions to identify potential pathways regulated by SARS-CoV-2. Briefly, HPAEpiC cells were reverse transfected with firefly luciferase reporter constructs containing response elements for the indicated pathways, and control Renilla luciferase constructs for 24 h. Cells were then pretreated with colchicine for 2 h and incubated with SARS-CoV-2 for 24 h. Then, the luciferase activities of the cells were measured with a dual-luciferase reporter assay system (E1910, Promega) on a fluorescent microplate reader (Molecular Devices Inc). Reporter luciferase activity was normalized to Renilla luciferase activity for each sample. All experiments were performed with three biological replicates.

### Immunofluorescence

Cells were fixed with 4% paraformaldehyde (PFA) for 10 min at room temperature. Paraformaldehyde-fixed, paraffin-embedded tissue was cut into 4 μm and adhered to frosted glass slides. After washed with PBS and treated with PBS containing 0.1% Triton X-100 for 15 min, the sections or cells were then permeabilized and blocked with PBST containing 5% fetal bovine serum for 90 min at room temperature. Cells were immunostained with anti-ACE2 (ab15348, 1:500 dilution, Abcam), or anti-Sp1 (T453) (ab59257, 1:500 dilution, Abcam), or anti-HNF4α antibodies (3113, 1:1000 dilution, Cell Signaling Technology) overnight at 4 °C. Tissue sections were immunostained with anti-ACE2 (GB11267, 1:200, Servicebio, Wuhan, China) or anti-SARS-CoV-2-N antibodies (40143-MM05, 1:500 dilution, SinoBiological, Beijing, China) overnight at 4°C. After washed three times with 0.1% Tween-20 in PBS (PBST), these cells were incubated with Alexa Fluor 594 anti-Rabbit IgG (H+L) (A-21207, 1:200 dilution, ThermoFisher Scientific), or Cy3 conjugated Goat anti-mouse IgG (H+L) (GB21301, 1:300, Servicebio), or Alexa Fluor 488-conjugated Goat anti-Rabbit IgG (H+L) (GB25303, 1:500, Servicebio) for 1 h. After staining with primary antibodies, nuclei were counterstained with DAPI. Images were acquired using a Zeiss Axioskop 2 plus fluorescence microscope (Carl Zeiss, Jena, Germany).

### Luciferase reporter assay

HPAEpiC cells at a density of 3×10^3^ were co-transfected with pRL-SV40 vector and phACE2- promoter-TA-luc (D2488, Beyotime) using Lipofectamine 3000 reagent (L3000015, Invitrogen). After 48 h of transfection, the luciferase activity was measured using the dual-luciferase reporter assay system (E1910, Promega, Shanghai, China) on a fluorescent microplate reader (Molecular Devices Inc., Sunnyvale, CA). The ratio of firefly luciferase to Renilla luciferase was calculated for each experiment and averaged from 3 replicates. All experiments were performed with three biological replicates.

### Transcription factor binding motif enrichment analysis

The ACE2 promoter sequence (1500 bp upstream of the transcription start site) was extracted from the NCBI database. The DNA-binding motifs of transcription factors ER, GATA6, HNF4α, NF-κB, and Sp1 were obtained from the JASPAR CORE database (JASPAR-A database of transcription factor binding profiles (genereg.net)). Then, the MEME Suite (Introduction- MEME Suite (meme-suite.org)) was used to interrogate the enrichment of these transcription factor motifs and the binding sites in the ACE2 promoter sequence (Bailey et al., 2015). FIMO analysis was performed using stringent criteria including *P*-value < 1E-4 and a maximum of two mismatched residues.

### ChIP-qPCR

ChIP was performed as described previously (Tao et al., 2016). Briefly, chromatin immunoprecipitation (ChIP) assay was performed using the ChIP assay kit (P2078, Beyotime) following the manufacturer’s directions as described. After crosslinking with formaldehyde, the chromatin solutions were sonicated and incubated with anti-Sp1 (9389, 1:100 dilution, Cell Signaling Technology), anti-HNF4α (ab181604, 1:100 dilution, Abcam) antibodies, and control IgG, and rotated overnight at 4 °C, respectively. After purified by a DNA purification kit (BioTeke Corp.), the immunoprecipitated DNA was detected for PCR analysis. All ChIP-qPCR experiments were performed with three biological replicates.

### RNA interference for cells

All chemically synthesized siRNAs were obtained from Gene-Pharma Corporation (Shanghai, China). To silence the expression of HNF4α, or Sp1, or AKT by siRNA, HPAEpiC were seeded at density of 5 × 10^5^ cells per well in 6-well plates containing complete culture medium. After 24 h, cells were transiently transfected with 100 nM of siRNAs using Lipofectamine 3000 (L3000015, Invitrogen, Beijing, China). Gene silencing efficiency was confirmed by qPCR 48 h post-transfection. The following siRNAs were used (sequence of the sense strand): HNF4α, 5’-GUC AUC GUU GCC AAC ACA AUG-3’; Sp1, 5’-CUC CAA GGC CUG GCU AAU AAU-3’; AKT, 5’-CGC GUG ACC AUG AAC GAG UUU-3’; negative control, 5’-UUC UCC GAA CGU GUC ACG UUU-3’.

### Co-immunoprecipitation

For co-immunoprecipitation experiments, HPAEpiC cells were lysed on ice for 30 min in cell lysis buffer (P0013, Beyotime, Shanghai, China). After centrifugation at 12000 rpm for 30 min at 4 °C. The supernatant was collected and incubated with anti-HNF4α antibodies (ab181604, 1:70 dilution, Abcam, Shanghai, China) overnight. After 4 h incubation with Protein AAgarose (20333, Thermo Scientific, Shanghai, China) at 4 °C, the complexes were washed three times. Immunoblotting was performed after elution.

### Western blot

For measurement of protein expression, HPAEpiC cells were re-suspended in RIPA buffer (R0278, Sigma-Aldrich, Shanghai, China) on ice for 1 hour. Protein samples were separated on SDS-PAGE gels, and then transferred to polyvinylidene fluoride (PVDF) membranes. After blocking with 5% BSA in PBS-T buffer containing 0.05% Tween-20, the membranes were incubated with primary antibodies overnight at 4 °C. Primary antibodies used in this study included: anti-ACE2 (ab108252, 1:1000 dilution, Abcam), anti-HNF4α (ab181604, 1:1000 dilution, Abcam), anti-phospho-AKT (Thr308) (13038, 1:1000 dilution, Cell Signaling Technology, Shanghai, China), anti-phospho-AKT (Ser473) (4060, 1:1000 dilution, Cell Signaling Technology), anti-pan-AKT (4691, 1:1000 dilution, Cell Signaling Technology), anti-phospho-Sp1 (T453) (ab59257, 1:1000 dilution, Abcam), and anti-Actin antibodies (sc- 47778, 1:5000 dilution, Santa Cruz-Biotechnology, Shanghai, China). After washed with PBS- T, the membranes were incubated with horseradish peroxidase (HRP)-conjugated secondary antibodies for 2 h at room temperature. The secondary antibodies used in our experiments were HRP-conjugated anti-mouse (7076, 1:2000 dilution, Cell Signaling Technology) or anti-rabbit IgG antibodies (7074, 1:2000 dilution, Cell Signaling Technology). The protein bands were detected using ECL (RPN2232, GE Healthcare, Little Chalfont, UK) on Amersham Imager 600. Subsequent image analysis was performed using ImageJ software. All experiments were performed with three biological replicates.

### Animal experiments and in vivo procedures

Syrian hamsters (*Mesocricetus auratus*) were purchased from Beijing Vital River Laboratory Animal Technology Co., Ltd. weighting 85-100 g, aged 5 weeks, male. In this study, all animals used were chosen randomly. The colchicine (C804812, Macklin, Shanghai, China) and MithA (A600668, Sangon Biotech, Shanghai, China) were dissolved in 1% (v/v) DMSO and 99% saline. The Syrian hamsters were anesthetized by inhalation of isoflurane and infected with 10^4^ TCID_50_ of SARS-CoV-2 by intranasal instillation. One hour later, 0.2 mg/kg of either colchicine or MithA were given i.p. once daily. Trachea, lung, and kidney tissues were collected on days 3 post infection.

### H&E staining and histopathology score

Lungs, trachea and kidney from Syrian hamsters were collected, and fixed with 4% paraformaldehyde and then paraffin-embedded. Tissue sections at 4 μm thickness were stained with hematoxylin and eosin (H&E) for histopathological examination. After H&E staining, a four-point scoring system was applied to assess the severity of pathology in tissues. The grades are expressed as numbers, where 0 indicates no pathological change and 1-4 indicate increasing severity. Lung histopathological scores were assessed and summarized based on alveolar wall thickening, edema, hemorrhage, and inflammatory cell infiltration. Kidney histopathological scores were assessed and summarized based on cellular degeneration, necrosis, hemorrhage, inflammatory cell infiltration, and congestion. Histopathological scores represent the sum of the injury subtype scores for each condition on a scale of 0-20.

### Statistical analysis

Differences in gene expression, mRNA and protein levels, Viral RNA, Luciferase reporter assay, ChIP-qPCR assay, and fluorescence intensity were assessed by performing Student’s t-test. Data were analyzed using GraphPad Prism 8 (GraphPad Software Inc., La Jolla, CA).

### Study approval

In this study, all animal experimental procedures were approved by the Institutional Committee for Animal Care and Biosafety at Kunming Institute of Zoology, Chinese Academy of Sciences (Permit Number: IACUC-RE-2021-10-002).

## Data availability

All data generated or analysed during this study are included in the manuscript and supporting file.

## Author contributions

C.G. Z., Y.T. Z., Y.C. M., H. H., and R.H. L. designed the experiments and analyzed the data. H. H., R.H. L., X.Y. L., L.Q. W., Q. Z., X.Y. T., and R. Z. D performed the experiments. C.G. Z., Y.T. Z., Y.C. M., H. H., and R.H. L. interpreted the data. C.G. Z., Y.T. Z., Y.C. M., and H. H. wrote the manuscript.

## Acknowledgements

We thank Dr. Changwen Ke (Guangdong Provincial Center for Disease Control and Prevention) for providing SARS-CoV-2 strain. We appreciate all the support from Kunming National High-level Biosafety Research Center for Non-human Primates, Kunming Institute of Zoology, Chinese Academy of Sciences. This work was supported in part by grants from the National Key Research and Development Program of China (2021YFC2301303), Yunnan Key Research and Development Program (202103AC100005, 202103AQ100001, 202102AA310055), and the Major Science and Technology Project in Yunnan Province of China (202001BB050001).

---

**Figure 1-figure supplement 1.**
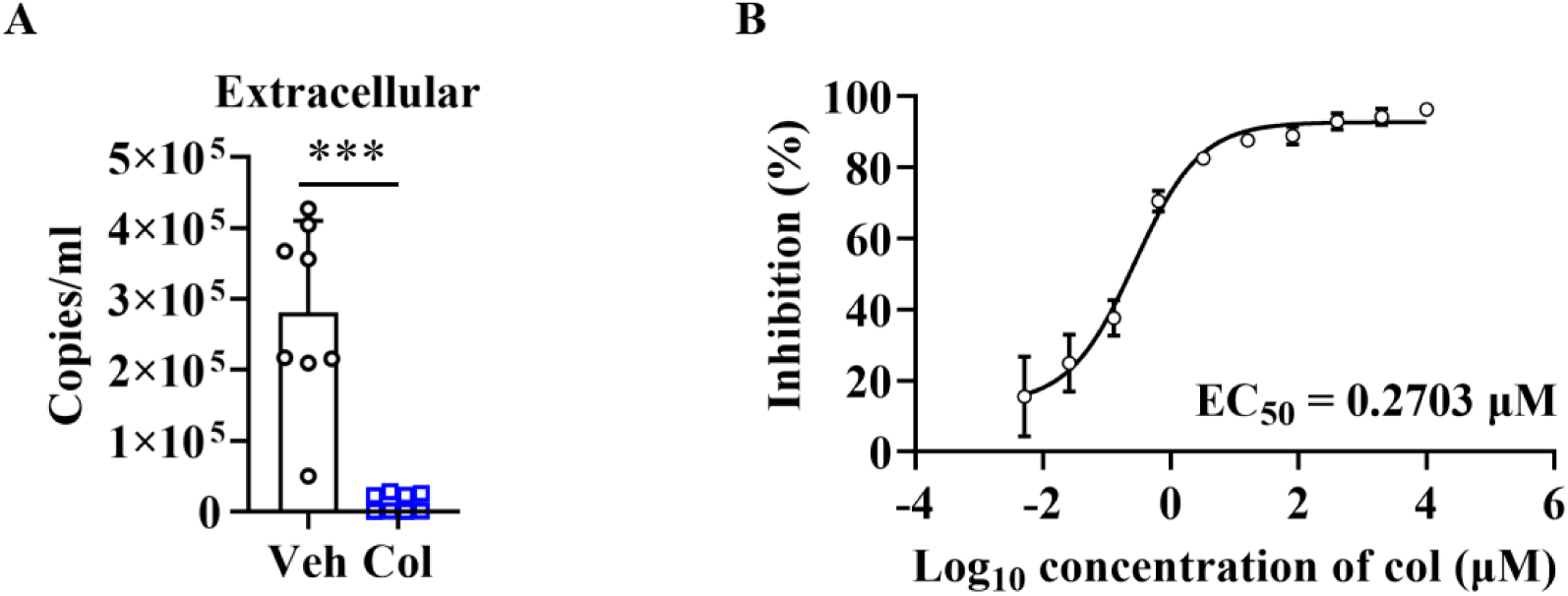
Colchicine blocks SARS-CoV-2 replication. (**A**) Colchicine (20 nM) inhibited SARS-CoV-2 replication in HPAEpiC cells. Cell culture supernatants were collected for viral load determination (n=8 each group). Error bars show means ± SD. \*\*\**P* < 0.001 (unpaired Student’s t test). (**B**) Dose-response analysis of HPAEpiC cells treated with colchicine at the indicated concentrations and infected with SARS-CoV-2 (MOI = 1) for 48 hours. **Figure 1-figure supplement 1-source data 1** Original file for determination of viral load in Figure 1-figure supplement 1A. **Figure 1-figure supplement 1-source data 2** Original file for dose-response analysis in Figure 1-figure supplement 1B.

**Figure 2-figure supplement 1.**
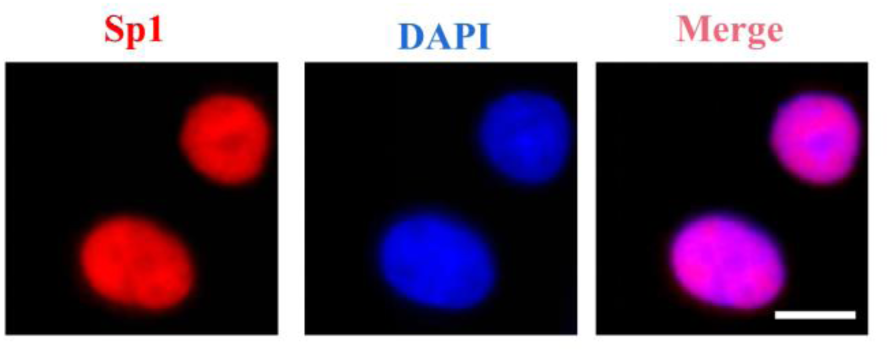
Sp1 is mainly located in nucleus of HPAEpiC cells.

**Figure 3-figure supplement 1.**
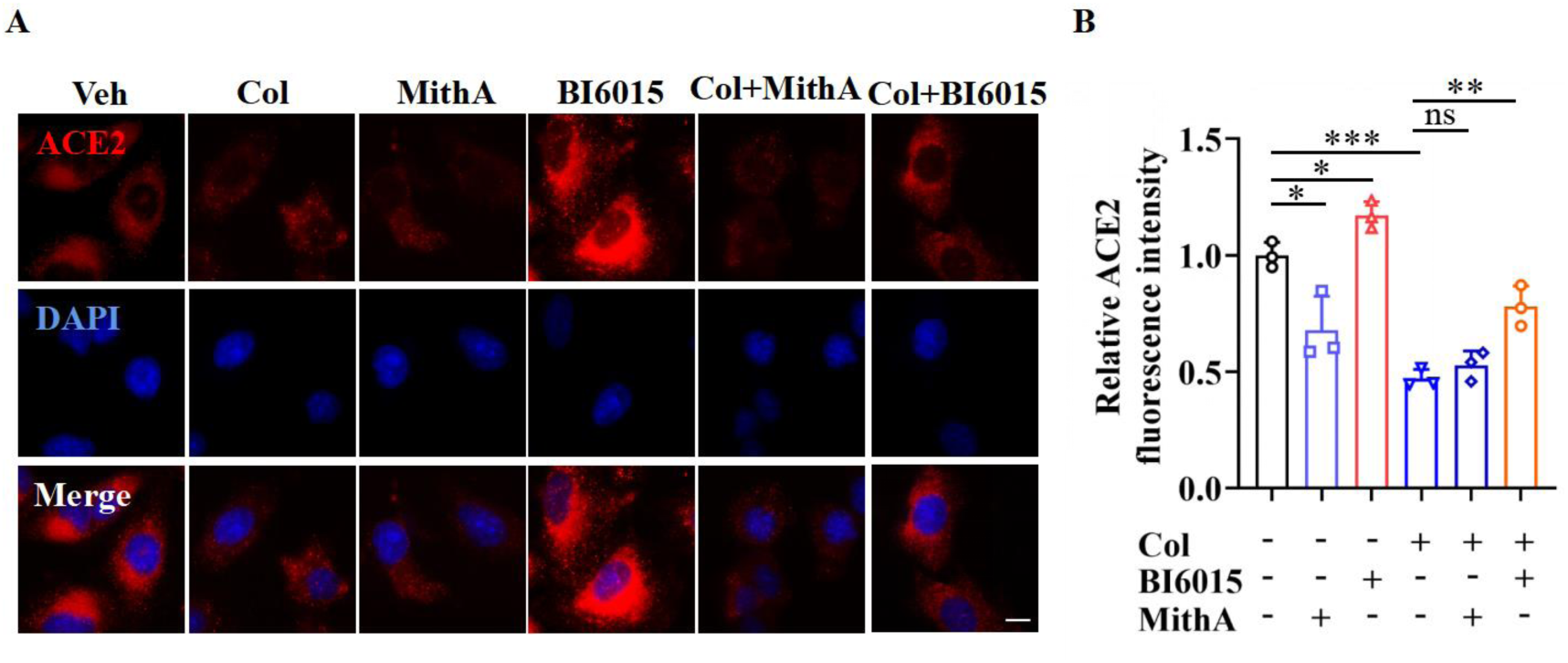
Immunofluorescence analysis of the expression and localization of ACE2 in A549 cells. (A) Representative images of immunofluorescence staining for ACE2 in A549 cells. A549 cells were treated with MithA, BI6015, colchicine, colchicine + MithA, or colchicine + BI6015. Scale bar: 10 μm. (**B**) Quantification of ACE2 fluorescence intensity. These results are means ± SD of three independent experiments. \**P* < 0.05, \*\**P* < 0.01, \*\*\**P* < 0.001, ns, not significant (unpaired Student’s t test). Veh, Vehicle. Col, Colchicine. **Figure 3-figure supplement 1-source data 1** Original file for quantification of ACE2 fluorescence intensity in Figure 3-figure supplement 1B.

**Figure 3-figure supplement 2.**
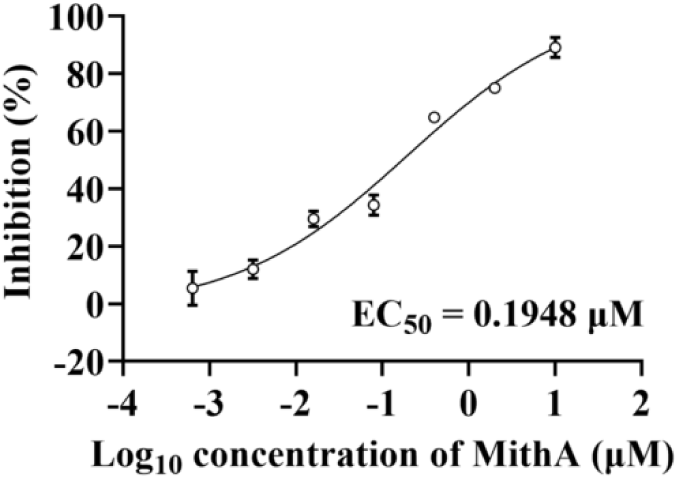
MithA inhibited SARS-CoV-2 replication in a dose- dependent manner. Dose-response analysis of HPAEpiC cells treated with the indicated concentrations of MithA and infected with SARS-CoV-2 (MOI = 1) for 48 hours. EC50 was achieved by plaque reduction assay and plotted using logistic non-linear regression model. **Figure 3-figure supplement 2-source data 1** Original file for dose-response analysis in Figure 3-figure supplement 2.

**Figure 5-figure supplement 1.**
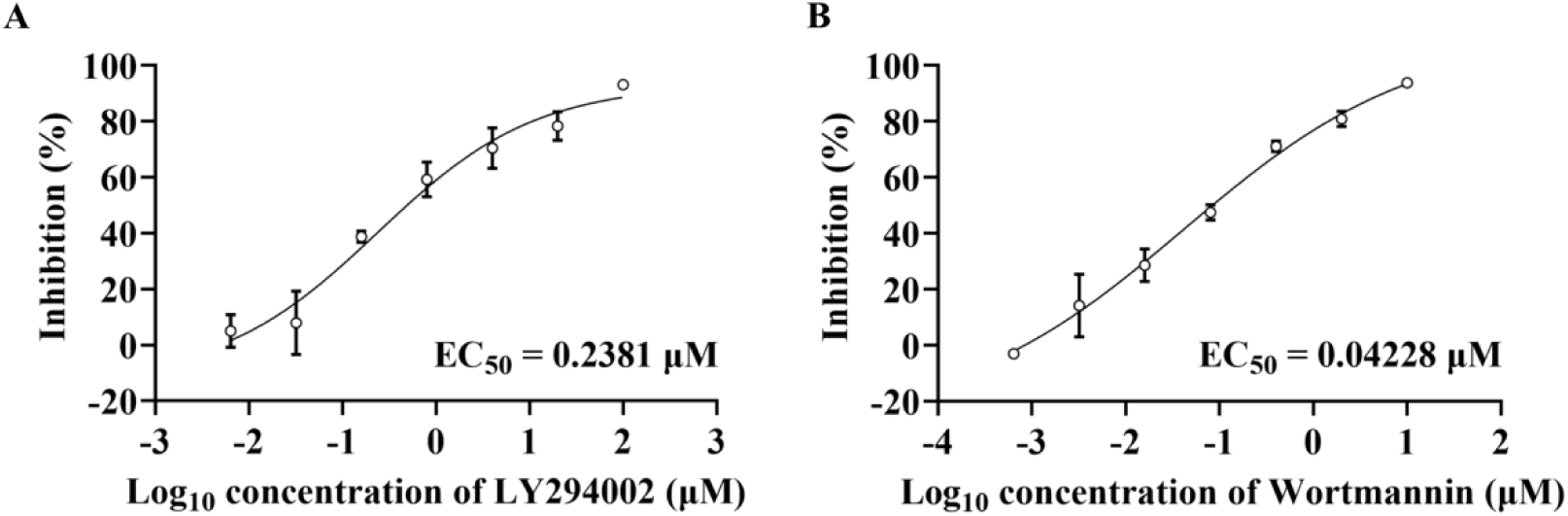
PI3K inhibitors inhibit SARS-CoV-2 replication in a dose- dependent manner. (**A and B**) Dose-response analysis of HPAEpiC cells treated with the indicated concentrations of PI3K/AKT inhibitors LY294002 (**A**) or wortmannin (**B**) and infected with SARS-CoV-2 (MOI = 1) for 48 hours. EC50 was achieved by plaque reduction assay and plotted using logistic non-linear regression model. **Figure 5-figure supplement 1-source data 1** Original file for dose-response analysis in Figure 5-figure supplement 1A and 1B.

**Figure 6-figure supplement 1.**
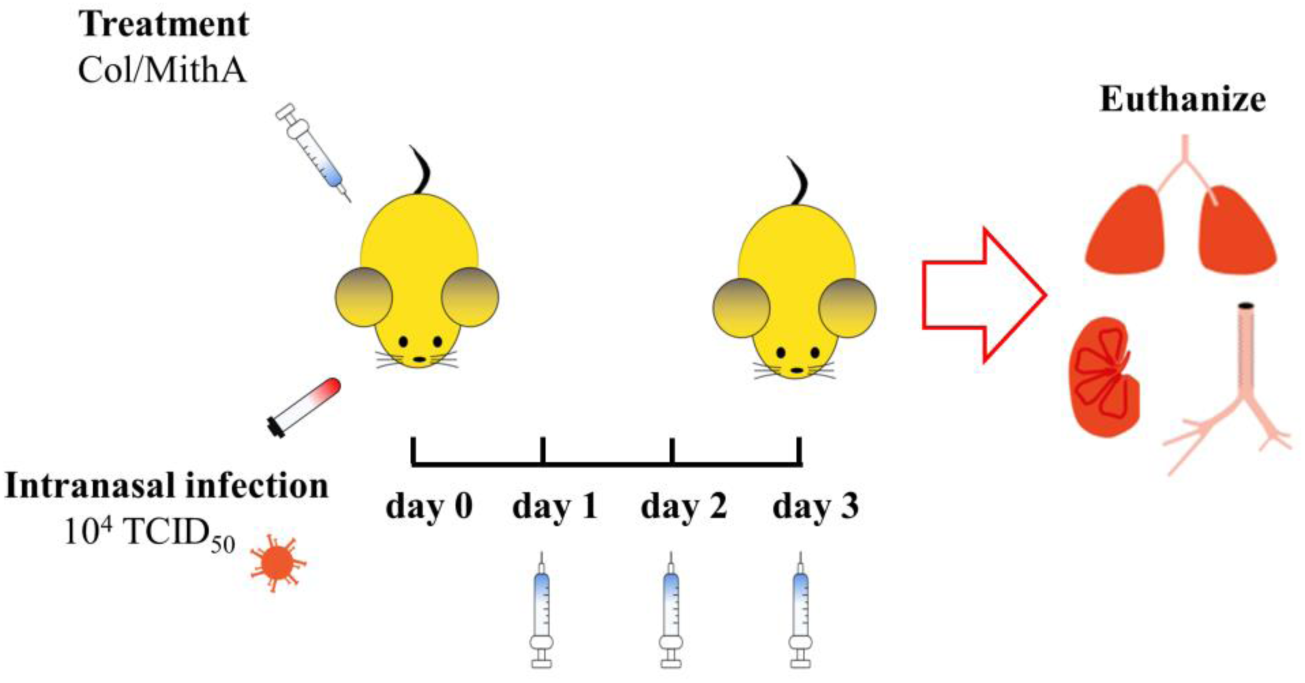
A schematic presentation of the experiment design for SARS-CoV-2 infection in Syrian hamsters. On day 0 of SARS-CoV-2 infection in Syrian hamsters, colchicine and MithA were injected intraperitoneally, respectively, at a dose of 0.2 mg/kg/day. According to different groups, the drug was injected up to the 3th day after infection. The uninfected and placebo groups were given the same dose of vehicle daily. Five animals of each group were euthanized at 3 days post infection.

**Figure 6-figure supplement 2.**
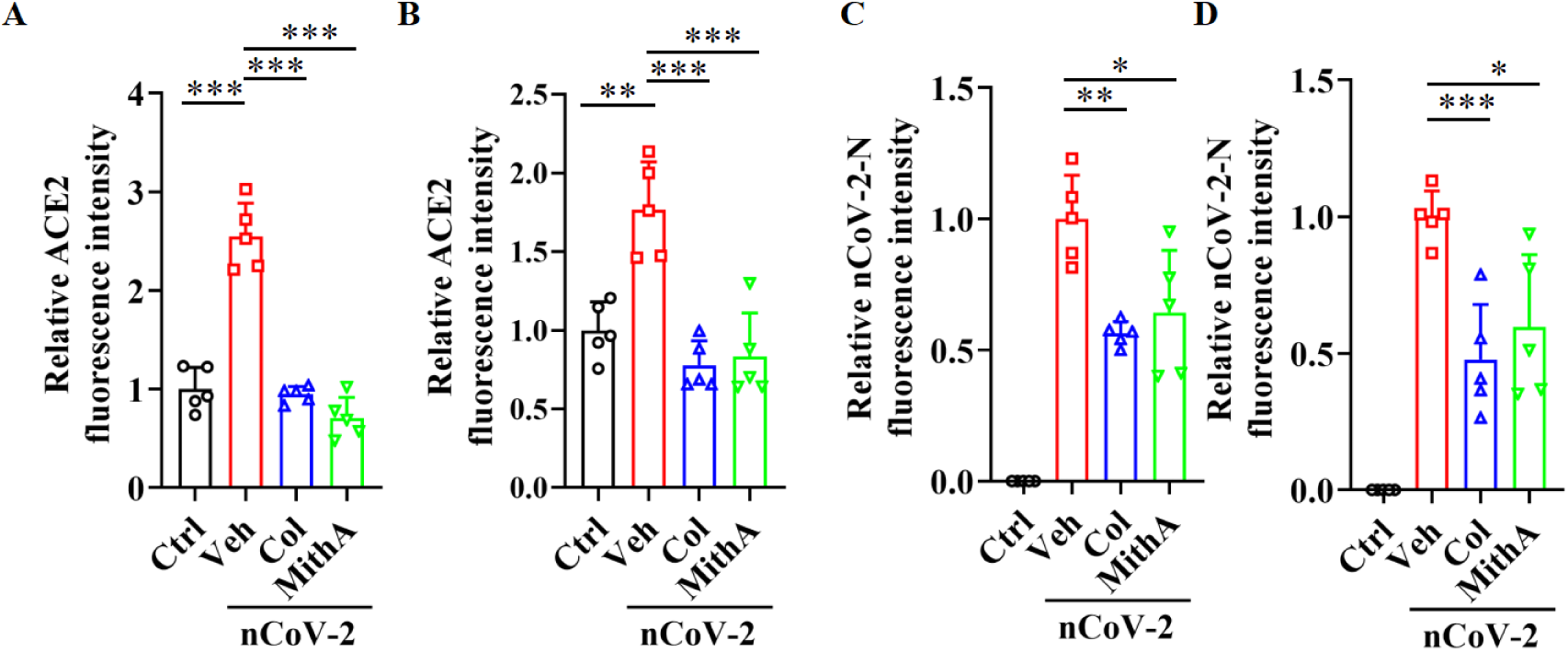
Either colchicine or MithA blocks the replication of SARS- CoV-2 by inhibiting the expression of ACE2 in both of the lung and trachea of hamsters. (A and B) Quantification of ACE2 fluorescence intensity in both of the lung (A) and trachea (B) of hamsters. Error bars show means ± SD. \*\**P* < 0.01, \*\*\**P* < 0.001 (Unpaired Student’ s t test). (**C and D**) Quantification of SARS-CoV-2-N fluorescence intensity in both of the lung (C) and trachea (**D**) of hamsters. Error bars show means ± SD. \**P* < 0.05, \*\**P* < 0.01, \*\*\**P* < 0.001 (Unpaired Student’s t test). Ctrl, Control. Veh, Vehicle. Col, Colchicine. nCoV-2, SARS- CoV-2. **Figure 6-figure supplement 2-source data 1** Original file for quantification of ACE2 fluorescence intensity in Figure 6-figure supplement 2A and 2B. **Figure 6-figure supplement 2-source data 2** Original file for quantification of SARS-CoV-2-N fluorescence intensity in Figure 6-figure supplement 2C and 2D.

## References

Adapala, N. S., Root, S., Lorenzo, J., Aguila, H. & Sanjay, A. 2019. PI3K activation increases SDF-1 production and number of osteoclast precursors, and enhances SDF-1- mediated osteoclast precursor migration. Bone Reports, 10, 100203.

Bailey, T. L., Johnson, J., Grant, C. E. & Noble, W. S. 2015. The MEME Suite. Nucleic Acids Research, 43, W39–49.

Brevini, T., Maes, M., Webb, G. J., John, B. V., Fuchs, C. D., Buescher, G., Wang, L., Griffiths, C., Brown, M. L., Scott, W. E., Pereyra-Gerber, P., Gelson, W. T. H., Brown, S., Dillon, S., Muraro, D., Sharp, J., Neary, M., Box, H., Tatham, L., Stewart, J., Curley, P., Pertinez, H., Forrest, S., Mlcochova, P., Varankar, S. S., Darvish-Damavandi, M., Mulcahy, V. L., Kuc, R. E., Williams, T. L., Heslop, J. A., Rossetti, D., Tysoe, O. C., Galanakis, V., Vila-Gonzalez, M., Crozier, T. W. M., Bargehr, J., Sinha, S., Upponi, S. S., Fear, C., Swift, L., Saeb-Parsy, K., Davies, S. E., Wester, A., Hagström, H., Melum, E., Clements, D., Humphreys, P., Herriott, J., Kijak, E., Cox, H., Bramwell, C., Valentijn, A., Illingworth, C. J. R., Dahman, B., Bastaich, D. R., Ferreira, R. D., Marjot, T., Barnes, E., Moon, A. M., Barritt, A. S., Gupta, R. K., Baker, S., Davenport, A. P., Corbett, G., Gorgoulis, V. G., Buczacki, S. J. A., Lee, J.- H., Matheson, N. J., Trauner, M., Fisher, A. J., Gibbs, P., Butler, A. J., Watson, C. J. E., Mells, G. F., Dougan, G., Owen, A., Lohse, A. W., Vallier, L., Sampaziotis, F. & Consortium, U.-P. R. 2022. FXR inhibition may protect from SARS- CoV-2 infection by reducing ACE2. Nature.

Callahan, V., Hawks, S., Crawford, M. A., Lehman, C. W., Morrison, H. A., Ivester, H. M., Akhrymuk, I., Boghdeh, N., Flor, R., Finkielstein, C. V., Allen, I. C., Weger-Lucarelli, J., Duggal, N., Hughes, M. A. & Kehn-Hall, K. 2021. The Pro-Inflammatory Chemokines CXCL9, CXCL10 and CXCL11 Are Upregulated Following Sars-CoV-2 Infection in an Akt-Dependent Manner. Viruses, 13.

Chan, J. F.-W., Zhang, A. J., Yuan, S., Poon, V. K.-M., Chan, C. C.-S., Lee, A. C.-Y., Chan, W.-M., Fan, Z., Tsoi, H.-W., Wen, L., Liang, R., Cao, J., Chen, Y., Tang, K., Luo, C., Cai, J.-P., Kok, K.-H., Chu, H., Chan, K.-H., Sridhar, S., Chen, Z., Chen, H., To, K. K.-W. & Yuen, K.-Y. 2020. Simulation of the Clinical and Pathological Manifestations of Coronavirus Disease 2019 (COVID-19) in a Golden Syrian Hamster Model: Implications for Disease Pathogenesis and Transmissibility. Clinical Infectious Diseases, 71, 2428–2446.

Choi, E.-S., Nam, J.-S., Jung, J.-Y., Cho, N.-P. & Cho, S.-D. 2014. Modulation of specificity protein 1 by mithramycin A as a novel therapeutic strategy for cervical cancer. Scientific Reports, 4, 7162.

Choudhary, S., Kanevsky, I., Yildiz, S., Sellers, R. S., Swanson, K. A., Franks, T., Rathnasinghe, R., Munoz-Moreno, R., Jangra, S., Gonzalez, O., Meade, P., Coskran, T., Qian, J., Lanz, T. A., Johnson, J. G., Tierney, C. A., Smith, J. D., Tompkins, K., Illenberger, A., Corts, P., Ciolino, T., Dormitzer, P. R., Dick, E. J., Shivanna, V., Hall-Ursone, S., Cole, J., Kaushal, D., Fontenot, J. A., Martinez-Romero, C., Mcmahon, M., Krammer, F., Schotsaert, M. & García-Sastre, A. 2022. Modeling Sars-CoV-2: Comparative Pathology in Rhesus Macaque and Golden Syrian Hamster Models. Toxicologic Pathology, 01926233211072767.

Chu, J., Xing, C., Du, Y., Duan, T., Liu, S., Zhang, P., Cheng, C., Henley, J., Liu, X., Qian, C., Yin, B., Wang, H. Y. & Wang, R.-F. 2021. Pharmacological inhibition of fatty acid synthesis blocks Sars-CoV-2 replication. Nature Metabolism, 3, 1466–1475.

Dasgeb, B., Kornreich, D., Mcguinn, K., Okon, L., Brownell, I. & Sackett, D. L. 2018. Colchicine: an ancient drug with novel applications. British Journal of Dermatology, 178, e167–e167.

Dong, M., Zhang, J., Ma, X., Tan, J., Chen, L., Liu, S., Xin, Y. & Zhuang, L. 2020. ACE2, TMPRSS2 distribution and extrapulmonary organ injury in patients with COVID-19. Biomedicine & Pharmacotherapy, 131, 110678.

Drosos, A. A., Pelechas, E., Drossou, V. & Voulgari, P. V. 2022. Colchicine Against Sars-CoV-2 Infection: What is the Evidence? Rheumatology and Therapy, 9, 379–389.

Elshafei, M. N., El-Bardissy, A., Khalil, A., Danjuma, M., Mubasher, M., Abubeker, I. Y. & Mohamed, M. F. H. 2021. Colchicine use might be associated with lower mortality in COVID-19 patients: A meta-analysis. European Journal of Clinical Investigation, 51, e13645.

Gao, C.-C., Li, M., Deng, W., Ma, C.-H., Chen, Y.-S., Sun, Y.-Q., Du, T., Liu, Q.-L., Li, W.-J., Zhang, B., Sun, L., Liu, S.-M., Li, F., Qi, F., Qu, Y., Ge, X., Liu, J., Wang, P., Niu, Y., Liang, Z., Zhao, Y.-L., Huang, B., Peng, X.-Z., Yang, Y., Qin, C., Tong, W.-M. & Yang, Y.-G. 2022. Differential transcriptomic landscapes of multiple organs from Sars-CoV- 2 early infected rhesus macaques. Protein & Cell, 13, 920–939.

Gasparyan, A. Y., Ayvazyan, L., Yessirkepov, M. & Kitas, G. D. 2015. Colchicine as an anti-inflammatory and cardioprotective agent. Expert Opinion on Drug Metabolism & Toxicology, 11, 1781–1794.

Gomez-Villafuertes, R., Garcia-Huerta, P., Diaz-Hernandez, J. I. & Miras-Portugal, M. T. 2015. PI3K/Akt signaling pathway triggers P2X7 receptor expression as a pro-survival factor of neuroblastoma cells under limiting growth conditions. Scientific Reports, 5, 18417.

Ho, J. S. Y., Mok, B. W.-Y., Campisi, L., Jordan, T., Yildiz, S., Parameswaran, S., Wayman, J. A., Gaudreault, N. N., Meekins, D. A., Indran, S. V., Morozov, I., Trujillo, J. D., Fstkchyan, Y. S., Rathnasinghe, R., Zhu, Z., Zheng, S., Zhao, N., White, K., Ray-Jones, H., Malysheva, V., Thiecke, M. J., Lau, S.-Y., Liu, H., Zhang, A. J., Lee, A. C.-Y., Liu, W.-C., Jangra, S., Escalera, A., Aydillo, T., Melo, B. S., Guccione, E., Sebra, R., Shum, E., Bakker, J., Kaufman, D. A., Moreira, A. L., Carossino, M., Balasuriya, U. B. R., Byun, M., Albrecht, R. A., Schotsaert, M., Garcia-Sastre, A., Chanda, S. K., Miraldi, E. R., Jeyasekharan, A. D., Tenoever, B. R., Spivakov, M., Weirauch, M. T., Heinz, S., Chen, H., Benner, C., Richt, J. A. & Marazzi, I. 2021. TOP1 inhibition therapy protects against Sars-CoV-2-induced lethal inflammation. Cell, 184, 2618–2632.e17.

Hoffmann, M., Kleine-Weber, H., Schroeder, S., Krüger, N., Herrler, T., Erichsen, S., Schiergens, T. S., Herrler, G., Wu, N. H., Nitsche, A., Müller, M. A., Drosten, C. & Pöhlmann, S. 2020. Sars-CoV-2 Cell Entry Depends on ACE2 and TMPRSS2 and Is Blocked by a Clinically Proven Protease Inhibitor. Cell, 181, 271–280.e8.

Inde, Z., Croker, B., Yapp, C., Joshi, G. N., Spetz, J., Fraser, C., Qin, X. P., Xu, L., Deskin, B., Ghelfi, E., Webb, G., Carlin, A. F., Zhu, Y. F. P. P., Leibel, S. L., Garretson, A. F., Clark, A. E., Duran, J. M., Pretorius, V., Crotty-Alexander, L. E., Li, C. D., Lee, J. C., Sodhi, C., Hackam, D. J., Sun, X., Hata, A. N., Kobzik, L., Miller, J., Park, J. A., Brownfield, D., Jia, H. P. & Sarosiek, K. A. 2021. Age-dependent regulation of Sars-CoV-2 cell entry genes and cell death programs correlates with COVID-19 severity. Science Advances, 7.

Jansen, J., Reimer, K. C., Nagai, J. S., Varghese, F. S., Overheul, G. J., De Beer, M., Roverts, R., Daviran, D., Fermin, L. A. S., Willemsen, B., Beukenboom, M., Djudjaj, S., Von Stillfried, S., Van Eijk, L. E., Mastik, M., Bulthuis, M., Dunnen, W. D., Van Goor, H., Hillebrands, J.-L., Triana, S. H., Alexandrov, T., Timm, M.-C., Van Den Berge, B. T., Van Den Broek, M., Nlandu, Q., Heijnert, J., Bindels, E. M. J., Hoogenboezem, R. M., Mooren, F., Kuppe, C., Miesen, P., Grünberg, K., Ijzermans, T., Steenbergen, E. J., Czogalla, J., Schreuder, M. F., Sommerdijk, N., Akiva, A., Boor, P., Puelles, V. G., Floege, J., Huber, T. B., Achdout, H., Aimon, A., Bar-David, E., Barr, H., Ben-Shmuel, A., Bennett, J., Boby, M. L., Borden, B., Bowman, G. R., Brun, J., Bvnbs, S., Calmiano, M., Carbery, A., Cattermole, E., Chernychenko, E., Choder, J. D., Clyde, A., Coffland, J. E., Cohen, G., Cole, J., Contini, A., Cox, L., Cvitkovic, M., Dias, A., Donckers, K., Dotson, D. L., Douangamath, A., Duberstein, S., Dudgeon, T., Dunnett, L., Eastman, P. K., Erez, N., Eyermann, C. J., Fairhead, M., Fate, G., Fearon, D., Federov, O., Ferla, M., Fernandes, R. S., Ferrins, L., Foster, R., Foster, H., Gabizon, R., Garcia-Sastre, A., Gawriljuk, V. O., Gehrtz, P., Gileadi, C., Giroud, C., Glass, W. G., Glen, R., Itai, G., Godoy, A. S., Gorichko, M., Gorrie-Stone, T., Griffen, E. J., Hart, S. H., Heer, J., Henry, M., et al. 2022. Sars-CoV-2 infects the human kidney and drives fibrosis in kidney organoids. Cell Stem Cell, 29, 217–231.e8.

Kamel, W., Noerenberg, M., Cerikan, B., Chen, H., Järvelin, A. I., Kammoun, M., Lee, J. Y., Shuai, N., Garcia-Moreno, M., Andrejeva, A., Deery, M. J., Johnson, N., Neufeldt, C. J., Cortese, M., Knight, M. L., Lilley, K. S., Martinez, J., Davis, I., Bartenschlager, R., Mohammed, S. & Castello, A. 2021. Global analysis of protein-RNA interactions in Sars-CoV-2-infected cells reveals key regulators of infection. Molecular Cell, 81, 2851–2867.e7.

Kiselyuk, A., Lee, S.-H., Farber-Katz, S., Zhang, M., Athavankar, S., Cohen, T., Pinkerton, Anthony B., Ye, M., Bushway, P., Richardson, Adam D., Hostetler, Heather A., Rodriguez-Lee, M., Huang, L., Spangler, B., Smith, L., Higginbotham, J., Cashman, J., Freeze, H., Itkin-Ansari, P., Dawson, Marcia I., Schroeder, F., Cang, Y., Mercola, M. & Levine, F. 2012. HNF4α Antagonists Discovered by a High-Throughput Screen for Modulators of the Human Insulin Promoter. Chemistry & Biology, 19, 806–818.

Klann, K., Bojkova, D., Tascher, G., Ciesek, S., Münch, C. & Cinatl, J. 2020. Growth Factor Receptor Signaling Inhibition Prevents Sars-CoV-2 Replication. Molecular Cell, 80, 164–174.e4.

Kuhlmann, C., Mayer, C. K., Claassen, M., Maponga, T., Burgers, W. A., Keeton, R., Riou, C., Sutherland, A. D., Suliman, T., Shaw, M. L. & Preiser, W. 2022. Breakthrough infections with Sars-CoV-2 omicron despite mRNA vaccine booster dose. The Lancet, 399, 625–626.

Lee, K.-A., Chae, J.-I. & Shim, J.-H. 2012. Natural diterpenes from coffee, cafestol and kahweol induce apoptosis through regulation of specificity protein 1 expression in human malignant pleural mesothelioma. Journal of Biomedical Science, 19, 60.

Lee, Y. C., Oslund, K. L., Thai, P., Velichko, S., Fujisawa, T., Duong, T., Denison, M. S. & Wu, R. 2011. 2,3,7,8-Tetrachlorodibenzo-p-dioxin–Induced MUC5AC Expression. American Journal of Respiratory Cell and Molecular Biology, 45, 270–276.

Legrand, M., Bell, S., Forni, L., Joannidis, M., Koyner, J. L., Liu, K. & Cantaluppi, V. 2021. Pathophysiology of COVID-19-associated acute kidney injury. Nature Reviews Nephrology, 17, 751–764.

Li, M., Wang, Y., Xia, X., Mo, P., Xu, J., Yu, C. & Li, W. 2019. Steroid receptor coactivator 3 inhibits hepatitis B virus gene expression through activating Akt signaling to prevent HNF4alpha nuclear translocation. Cell & Bioscience, 9, 64.

Liu, C., Ginn, H. M., Dejnirattisai, W., Supasa, P., Wang, B., Tuekprakhon, A., Nutalai, R., Zhou, D., Mentzer, A. J., Zhao, Y., Duyvesteyn, H. M. E., López-Camacho, C., Slon-Campos, J., Walter, T. S., Skelly, D., Johnson, S. A., Ritter, T. G., Mason, C., Costa Clemens, S. A., Gomes Naveca, F., Nascimento, V., Nascimento, F., Fernandes Da Costa, C., Resende, P. C., Pauvolid-Correa, A., Siqueira, M. M., Dold, C., Temperton, N., Dong, T., Pollard, A. J., Knight, J. C., Crook, D., Lambe, T., Clutterbuck, E., Bibi, S., Flaxman, A., Bittaye, M., Belij-Rammerstorfer, S., Gilbert, S. C., Malik, T., Carroll, M. W., Klenerman, P., Barnes, E., Dunachie, S. J., Baillie, V., Serafin, N., Ditse, Z., DA Silva, K., Paterson, N. G., Williams, M. A., Hall, D. R., Madhi, S., Nunes, M. C., Goulder, P., Fry, E. E., Mongkolsapaya, J., Ren, J., Stuart, D. I. & Screaton, G. R. 2021. Reduced neutralization of Sars-CoV-2 B.1.617 by vaccine and convalescent serum. Cell, 184, 4220–4236.e13.

Lu, R., Zhao, X., Li, J., Niu, P., Yang, B., Wu, H., Wang, W., Song, H., Huang, B., Zhu, N., Bi, Y., Ma, X., Zhan, F., Wang, L., Hu, T., Zhou, H., Hu, Z., Zhou, W., Zhao, L., Chen, J., Meng, Y., Wang, J., Lin, Y., Yuan, J., Xie, Z., Ma, J., Liu, W. J., Wang, D., Xu, W., Holmes, E. C., Gao, G. F., Wu, G., Chen, W., Shi, W. & Tan, W. 2020. Genomic characterisation and epidemiology of 2019 novel coronavirus: implications for virus origins and receptor binding. The Lancet, 395, 565–574.

Lu, T., Wang, Y. & Guo, T. 2022. Multi-omics in COVID-19: Seeing the unseen but overlooked in the clinic. Cell Reports Medicine, 3, 100580.

Manzini, M. C., Xiong, L., Shaheen, R., Tambunan, Dimira E., Di Costanzo, S., Mitisalis, V., Tischfield, David J., Cinquino, A., Ghaziuddin, M., Christian, M., Jiang, Q., Laurent, S., Nanjiani, ZOHAIR A., Rasheed, S., Hill, R. S., Lizarraga, Sofia B., Gleason, D., Sabbagh, D., Salih, Mustafa A., Alkuraya, Fowzan S. & Walsh, Christopher A. 2014. CC2D1A Regulates Human Intellectual and Social Function as well as NF-κB Signaling Homeostasis. Cell Reports, 8, 647–655.

Mccallum, M., Czudnochowski, N., Rosen, L. E., Zepeda, S. K., Bowen, J. E., Walls, A. C., Hauser, K., Joshi, A., Stewart, C., Dillen, J. R., Powell, A. E., Croll, T. I., Nix, J., Virgin, H. W., Corti, D., Snell, G. & Veesler, D. 2022. Structural basis of Sars-CoV-2 Omicron immune evasion and receptor engagement. Science, 375, 864–868.

Milanini-Mongiat, J., Pouysségur, J. & Pagès, G. 2002. Identification of Two Sp1 Phosphorylation Sites for p42/p44 Mitogen-activated Protein Kinases: THEIR IMPLICATION IN VASCULAR ENDOTHELIAL GROWTH FACTOR GENE TRANSCRIPTION*. Journal of Biological Chemistry, 277, 20631–20639.

Monteil, V., Kwon, H., Prado, P., Hagelkrüys, A., Wimmer, R. A., Stahl, M., Leopoldi, A., Garreta, E., Hurtado Del Pozo, C., Prosper, F., Romero, J. P., Wirnsberger, G., Zhang, H., Slutsky, A. S., Conder, R., Montserrat, N., Mirazimi, A. & Penninger, J. M. 2020. Inhibition of Sars-CoV-2 Infections in Engineered Human Tissues Using Clinical-Grade Soluble Human ACE2. Cell, 181, 905–913.e7.

Muñoz-Fontela, C., Dowling, W. E., Funnell, S. G. P., Gsell, P.-S., Riveros-Balta, A. X., Albrecht, R. A., Andersen, H., Baric, R. S., Carroll, M. W., Cavaleri, M., Qin, C., Crozier, I., Dallmeier, K., De Waal, L., De Wit, E., Delang, L., Dohm, E., Duprex, W. P., Falzarano, D., Finch, C. L., Frieman, M. B., Graham, B. S., Gralinski, L. E., Guilfoyle, K., Haagmans, B. L., Hamilton, G. A., Hartman, A. L., Herfst, S., Kaptein, S. J. F., Klimstra, W. B., Knezevic, I., Krause, P. R., Kuhn, J. H., LE Grand, R., Lewis, M. G., Liu, W.-C., Maisonnasse, P., Mcelroy, A. K., Munster, V., Oreshkova, N., Rasmussen, A. L., Rocha-Pereira, J., Rockx, B., Rodríguez, E., Rogers, T. F., Salguero, F. J., Schotsaert, M., Stittelaar, K. J., Thibaut, H. J., Tseng, C.-T., Vergara-Alert, J., Beer, M., Brasel, T., Chan, J. F. W., García-Sastre, A., Neyts, J., Perlman, S., Reed, D. S., Richt, J. A., Roy, C. J., Segalés, J., Vasan, S. S., Henao-Restrepo, A. M. & Barouch, D. H. 2020. Animal models for COVID-19. Nature, 586, 509–515.

Nadim, M. K., Forni, L. G., Mehta, R. L., Connor, M. J., Liu, K. D., Ostermann, M., Rimmelé, T., Zarbock, A., Bell, S., Bihorac, A., Cantaluppi, V., Hoste, E., Husain-Syed, F., Germain, M. J., Goldstein, S. L., Gupta, S., Joannidis, M., Kashani, K., Koyner, J. L., Legrand, M., Lumlertgul, N., Mohan, S., Pannu, N., Peng, Z., Perez-Fernandez, X. L., Pickkers, P., Prowle, J., Reis, T., Srisawat, N., Tolwani, A., Vijayan, A., Villa, G., Yang, L., Ronco, C. & Kellum, J. A. 2020. COVID-19-associated acute kidney injury: consensus report of the 25th Acute Disease Quality Initiative (ADQI) Workgroup. Nature Reviews Nephrology, 16, 747–764.

Pan, Y., Du, J., Liu, J., Wu, H., Gui, F., Zhang, N., Deng, X., Song, G., Li, Y., Lu, J., Wu, X., Zhan, S., Jing, Z., Wang, J., Yang, Y., Liu, J., Chen, Y., Chen, Q., Zhang, H., Hu, H., Duan, K., Wang, M., Wang, Q. & Yang, X. 2021. Screening of potent neutralizing antibodies against Sars-CoV-2 using convalescent patients-derived phage-display libraries. Cell Discovery, 7, 57.

Qiao, Y. Y., Wang, X. M., Mannan, R., Pitchiaya, S., Zhang, Y. P., Wotring, J. W., Xiao, L. B., Robinson, D. R., Wu, Y. M., Tien, J. C. Y., Cao, X. H., Simko, S. A., Apel, I. J., Bawa, P., Kregel, S., Narayanan, S. P., Raskind, G., Ellison, S. J., Parolia, A., Zelenka-Wang, S., Mcmurry, L., Su, F. Y., Wang, R., Cheng, Y. H., Delekta, A. D., Mei, Z. J., Pretto, C. D., Wang, S. M., Mehra, R., Sexton, J. Z. & Chinnaiyan, A. M. 2021. Targeting transcriptional regulation of Sars-CoV-2 entry factors ACE2 and TMPRSS2. Proceedings of the National Academy of Sciences of the United States of America, 118.

Rosenke, K., Hansen, F., Schwarz, B., Feldmann, F., Haddock, E., Rosenke, R., Barbian, K., Meade-White, K., Okumura, A., Leventhal, S., Hawman, D. W., Ricotta, E., Bosio, C. M., Martens, C., Saturday, G., Feldmann, H. & Jarvis, M. A. 2021. Orally delivered MK-4482 inhibits Sars-CoV-2 replication in the Syrian hamster model. Nature Communications, 12, 2295.

Samuel, R. M., Majd, H., Richter, M. N., Ghazizadeh, Z., Zekavat, S. M., Navickas, A., Ramirez, J. T., Asgharian, H., Simoneau, C. R., Bonser, L. R., Koh, K. D., Garcia-Knight, M., Tassetto, M., Sunshine, S., Farahvashi, S., Kalantari, A., Liu, W., Andino, R., Zhao, H., Natarajan, P., Erle, D. J., Ott, M., Goodarzi, H. & Fattahi, F. 2020. Androgen Signaling Regulates Sars-CoV-2 Receptor Levels and Is Associated with Severe COVID-19 Symptoms in Men. Cell Stem Cell, 27, 876–889.e12.

Schlesinger, N., Firestein, B. L. & Brunetti, L. 2020. Colchicine in COVID-19: an Old Drug, New Use. Current Pharmacology Reports, 6, 137–145.

Sia, S. F., Yan, L.-M., Chin, A. W. H., Fung, K., Choy, K.-T., Wong, A. Y. L., Kaewpreedee, P., Perera, R. A. P. M., Poon, L. L. M., Nicholls, J. M., Peiris, M. & Yen, H.-L. 2020. Pathogenesis and transmission of Sars-CoV-2 in golden hamsters. Nature, 583, 834–838.

Slobodnick, A., Shah, B., Pillinger, M. H. & Krasnokutsky, S. 2015. Colchicine: Old and New. The American Journal of Medicine, 128, 461–470.

Smith, K. D. & Akilesh, S. 2021. Pathogenesis of coronavirus disease 2019-associated kidney injury. Current Opinion in Nephrology and Hypertension, 30.

Su, C.-H., Wang, C.-Y., Lan, K.-H., Li, C.-P., Chao, Y., Lin, H.-C., Lee, S.-D. & Lee, W.-P. 2011. Akt phosphorylation at Thr308 and Ser473 is required for CHIP-mediated ubiquitination of the kinase. Cellular Signalling, 23, 1824–1830.

Sun, F., Mu, C., Kwok, H. F., Xu, J., Wu, Y., Liu, W., Sabatier, J. M., Annweiler, C., Li, X., Cao, Z. & Xie, Y. 2021. Capivasertib restricts Sars-CoV-2 cellular entry: a potential clinical application for COVID-19. International Journal of Biological Sciences, 17, 2348–2355.

Tao, J., Ma, Y.-C., Yang, Z.-S., Zou, C.-G. & Zhang, K.-Q. 2016. Octopamine connects nutrient cues to lipid metabolism upon nutrient deprivation. Science Advances, 2, e1501372.

Van Eijk, L. E., Binkhorst, M., Bourgonje, A. R., Offringa, A. K., Mulder, D. J., Bos, E. M., Kolundzic, N., Abdulle, A. E., Van Der Voort, P. H. J., Olde Rikkert, M. G. M., Van Der Hoeven, J. G., DEN Dunnen, W. F. A., Hillebrands, J.-L. & Van Goor, H. 2021. COVID-19: immunopathology, pathophysiological mechanisms, and treatment options. The Journal of Pathology, 254, 307–331.

Verdecchia, P., Cavallini, C., Spanevello, A. & Angeli, F. 2020. The pivotal link between ACE2 deficiency and Sars-CoV-2 infection. European Journal of Internal Medicine, 76, 14–20.

Wei, J., Alfajaro, M. M., Deweirdt, P. C., Hanna, R. E., Lu-Culligan, W. J., Cai, W. L., Strine, M. S., Zhang, S.-M., Graziano, V. R., Schmitz, C. O., Chen, J. S., Mankowski, M. C., Filler, R. B., Ravindra, N. G., Gasque, V., De Miguel, F. J., Patil, A., Chen, H., Oguntuyo, K. Y., Abriola, L., Surovtseva, Y. V., Orchard, R. C., Lee, B., Lindenbach, B. D., Politi, K., Van Dijk, D., Kadoch, C., Simon, M. D., Yan, Q., Doench, J. G. & Wilen, C. B. 2021. Genome-wide CRISPR Screens Reveal Host Factors Critical for Sars-CoV-2 Infection. Cell, 184, 76–91.e13.

Wilhelm, A., Widera, M., Grikscheit, K., Toptan, T., Schenk, B., Pallas, C., Metzler, M., Kohmer, N., Hoehl, S., Marschalek, R., Herrmann, E., Helfritz, F. A., Wolf, T., Goetsch, U. & Ciesek, S. 2022. Limited neutralisation of the Sars-CoV-2 Omicron subvariants BA.1 and BA.2 by convalescent and vaccine serum and monoclonal antibodies. eBioMedicine, 82, 104158.

Wu, F., Zhao, S., Yu, B., Chen, Y.-M., Wang, W., Song, Z.-G., Hu, Y., Tao, Z.-W., Tian, J.-H., Pei, Y.-Y., Yuan, M.-L., Zhang, Y.-L., Dai, F.-H., Liu, Y., Wang, Q.-M., Zheng, J.-J., Xu, L., Holmes, E. C. & Zhang, Y.-Z. 2020. A new coronavirus associated with human respiratory disease in China. Nature, 579, 265–269.

Wysocki, J., Ye, M., Hassler, L., Gupta, A. K., Wang, Y., Nicoleascu, V., Randall, G., Wertheim, J. A. & Batlle, D. 2021. A Novel Soluble ACE2 Variant with Prolonged Duration of Action Neutralizes Sars-CoV-2 Infection in Human Kidney Organoids. Journal of the American Society of Nephrology, 32, 795.

Xu, G., Li, Y., Zhang, S., Peng, H., Wang, Y., Li, D., Jin, T., He, Z., Tong, Y., Qi, C., Wu, G., Dong, K., Gou, J., Liu, Y., Xiao, T., Qu, J., Li, L., Liu, L., Zhao, P., Zhang, Z. & Yuan, J. 2021a. Sars-CoV-2 promotes RIPK1 activation to facilitate viral propagation. Cell Research, 31, 1230–1243.

Xu, Y., Zhang, S., Liao, X., Li, M., Chen, S., Li, X., Wu, X., Yang, M., Tang, M., Hu, Y., Li, Z., Yu, R., Huang, M., Song, L. & Li, J. 2021b. Circular RNA circIKBKB promotes breast cancer bone metastasis through sustaining NF-κB/bone remodeling factors signaling. Molecular Cancer, 20, 98.

Yan, R., Zhang, Y., Li, Y., Xia, L., Guo, Y. & Zhou, Q. 2020. Structural basis for the recognition of Sars-CoV-2 by full-length human ACE2. Science, 367, 1444–1448.

Yin, W., Xu, Y., Xu, P., Cao, X., Wu, C., Gu, C., He, X., Wang, X., Huang, S., Yuan, Q., Wu, K., Hu, W., Huang, Z., Liu, J., Wang, Z., Jia, F., Xia, K., Liu, P., Wang, X., Song, B., Zheng, J., Jiang, H., Cheng, X., Jiang, Y., Deng, S.-J. & Xu, H. E. 2022. Structures of the Omicron spike trimer with ACE2 and an anti-Omicron antibody. Science, 375, 1048–1053.

Zhao, J., Ye, W., Wu, J., Liu, L., Yang, L., Gao, L., Chen, B., Zhang, F., Yang, H. & Li, Y. 2015. Sp1-CD147 positive feedback loop promotes the invasion ability of ovarian cancer. Oncology Reports, 34, 67–76.

Zhou, Y., Wang, M., Li, Y., Wang, P., Zhao, P., Yang, Z., Wang, S., Zhang, L., Li, Z., Jia, K., Zhong, C., Li, N., Yu, Y. & Hou, J. 2021. Sars-CoV-2 Spike protein enhances ACE2 expression via facilitating Interferon effects in bronchial epithelium. Immunology Letters, 237, 33–41.

Zhuang, M.-W., Cheng, Y., Zhang, J., Jiang, X.-M., Wang, L., Deng, J. & Wang, P.-H. 2020. Increasing host cellular receptor—angiotensin-converting enzyme 2 expression by coronavirus may facilitate 2019-nCoV (or Sars-CoV-2) infection. Journal of Medical Virology, 92, 2693–2701.

Ziegler, C. G. K., Allon, S. J., Nyquist, S. K., Mbano, I. M., Miao, V. N., Tzouanas, C. N., Cao, Y., Yousif, A. S., Bals, J., Hauser, B. M., Feldman, J., Muus, C., Wadsworth, M. H., Kazer, S. W., Hughes, T. K., Doran, B., Gatter, G. J., Vukovic, M., Taliaferro, F., Mead, B. E., Guo, Z., Wang, J. P., Gras, D., Plaisant, M., Ansari, M., Angelidis, I., Adler, H., Sucre, J. M. S., Taylor, C. J., Lin, B., Waghray, A., Mitsialis, V., Dwyer, D. F., Buchheit, K. M., Boyce, J. A., Barrett, N. A., Laidlaw, T. M., Carroll, S. L., Colonna, L., Tkachev, V., Peterson, C. W., Yu, A., Zheng, H. B., Gideon, H. P., Winchell, C. G., Lin, P. L., Bingle, C. D., Snapper, S. B., Kropski, J. A., Theis, F. J., Schiller, H. B., Zaragosi, L.-E., Barbry, P., Leslie, A., Kiem, H.-P., Flynn, J. L., Fortune, S. M., Berger, B., Finberg, R. W., Kean, L. S., Garber, M., Schmidt, A. G., Lingwood, D., Shalek, A. K., Ordovas-Montanes, J., Banovich, N., Barbry, P., Brazma, A., Desai, T., Duong, T. E., Eickelberg, O., Falk, C., Farzan, M., Glass, I., Haniffa, M., Horvath, P., Hung, D., Kaminski, N., Krasnow, M., Kropski, J. A., Kuhnemund, M., Lafyatis, R., Lee, H., Leroy, S., Linnarson, S., Lundeberg, J., Meyer, K., Misharin, A., Nawijn, M., Nikolic, M. Z., Ordovas-Montanes, J., Pe’er, D., Powell, J., Quake, S., Rajagopal, J., Tata, P. R., Rawlins, E. L., Regev, A., Reyfman, P. A., Rojas, M., et al. 2020. Sars-CoV-2 Receptor ACE2 Is an Interferon-Stimulated Gene in Human Airway Epithelial Cells and Is Detected in Specific Cell Subsets across Tissues. Cell, 181, 1016–1035.e19.

